# The histidine kinase NahK in *Pseudomonas aeruginosa* is essential for nitric oxide-stress resistance in nutrient starved media

**DOI:** 10.1101/2025.07.03.662950

**Authors:** Sweta Anantharaman, Jiayuan Fu, Elizabeth M. Boon

**Affiliations:** Department of Chemistry, Stony Brook University, Stony Brook, NY, USA; Institute of Chemical Biology and Drug Discovery, Stony Brook University, Stony Brook, NY, USA

## Abstract

Biofilms are communities of bacteria growing within a matrix composed of polymeric substances that act as a barrier to antimicrobials, making bacteria within a biofilm recalcitrant to conventional antibiotic treatments. This lifestyle of bacteria is especially relevant during its pathogenesis, as observed in the case of *Pseudomonas aeruginosa* infections of immunocompromised patients. However, pico- to nanomolar concentrations of nitric oxide (NO) have been shown to be efficient in triggering *P. aeruginosa* to disperse from biofilms, suggesting that a combination of NO exposure and antibiotic treatment may help in mitigating infections. In *P. aeruginosa*, the NosP-NahK two component system, which comprises a NO sensing hemoprotein, NosP, and its associated hybrid histidine kinase, NahK, has been proven to be necessary for NO mediated biofilm dispersal. NahK also has additional roles in biofilm formation, motility, denitrification and virulence, due to its regulation of a global post-transcriptional regulator RsmA. Here, we uncover a novel role of NahK in enhancing the resistance of *P. aeruginosa* to signaling concentrations of NO. Deletion of *nahK* sensitizes the strain to nanomolar levels of NO, resulting in DNA damage. Consequently, the SOS stress response pathway is induced, which causes phenotypes such as cell filamentation and cell clustering due to lysis-mediated release of extracellular DNA. Our data also indicate that increased susceptibility of the Δ*nahK* strain to NO is due to the antagonization of RsmA, and is restricted to amino acid starved media, suggesting that NahK may also have previously unappreciated roles in modulating amino acid metabolism.

**Importance:** *P. aeruginosa* infections of cystic fibrosis patients can result in increased risk of fatality, necessitating the development of better therapeutic strategies to treat its infections. Here, we present data suggesting that nanomolar concentrations of NO, that are normally used to disperse *P. aeruginosa* from biofilms, can also sensitize the bacterium to NO mediated DNA damage upon loss of function of the histidine kinase NahK. The resulting SOS stress response may benefit the bacterium through favorable mutations but also poses a risk due to SOS-associated pyocin production through cell lysis. Hence, understanding the molecular mechanisms underlying NahK mediated resistance to NO is important to leverage this regulation and identify novel targets to treat *P. aeruginosa* infections.

## Introduction

The opportunistic pathogen *Pseudomonas aeruginosa* is ubiquitous in nature and is the leading cause of hospital acquired infections in immunocompromised patients (1, 2). This includes infections of burn wounds, catheters, urinary tracts and medical implants, and nosocomial infections such as those in the lungs of patients with cystic fibrosis. *P. aeruginosa* infection of cystic fibrosis patients is chronic and especially concerning as it increases the risk of respiratory failure and mortality in these patients (3–5). Hence, understanding its pathogenesis and mitigating its associated infections is of utmost importance. One of the main reasons why *P. aeruginosa* infections are difficult to treat is the formation of biofilms, which are communities of bacteria growing within a self-secreted polymeric matrix (6). This polymeric matrix is composed of extracellular polymeric substances (EPS) such as proteins, lipids, poly-uronic acids, and DNA, which makes penetration of chemicals such as antibiotics very difficult. Because of this, bacteria within a biofilm are 10-1000 times more resistant to conventional antibiotic treatments (7–9). Bacteria within the biofilm are sessile and survive inside until they exhaust all nutrients, following which they regain motility, disperse and attach to new surfaces that offer nutrients for their survival. In *P. aeruginosa*, this switch between sessile and motile lifestyles is regulated by the protein RsmA (10–12).

RsmA is a global post-transcriptional regulatory protein that affects the translation of more than 500 genes in *P. aeruginosa* (12, 13) . It recognizes GGA motifs on the hexaloops of its target mRNAs and binds to them, thereby preventing translation of the mRNA. This function of RsmA is referred to as its ‘ON’ state and can be turned ‘OFF’ by the small regulatory RNAs *rsmY* and *rsmZ* which block RsmA’s inhibitory function by binding to it through their GGA motifs (13). The switch in function of RsmA is regulated by the Gac multikinase network (MKN) comprising five kinases; GacS, RetS, SagS, PA1611 and NahK. These kinases regulate the sessile-to-motile lifestyle switch in *P. aeruginosa* through RsmA by modulating *rsmY*/*Z* transcription in response to environmental cues (14). Recent studies have shown that of these five kinases, NahK is one of the main regulators of RsmA’s function and does so by affecting *rsmY* and *rsmZ* transcription in a cyclic dimeric guanosine monophosphate (cyclic-di-GMP) dependent manner (15). When the kinase activity of NahK is inhibited, it decreases the phosphoryl flux through a phosphotransfer protein HptB, and unphosphorylated HptB promotes the intracellular accumulation of cyclic-di-GMP by activation of the diguanylate cyclases (DGC) HsbA, WspR, and a third DGC that is currently unknown. Increased levels of cyclic-di-GMP, which act as a secondary messenger molecule inside the cell, activates the transcription of *rsmY*, which turns RsmA OFF. When RsmA is OFF, *P. aeruginosa* favors sessile or biofilm lifestyle through de-repressing the translation of genes responsible for the formation of biofilms. We have recently shown that deletion of *nahK* from the *P. aeruginosa* PA14 genome promotes biofilm formation, decreases swarming motility, and increases the secretion of an exotoxin pyocyanin, which is consistent with the RsmA OFF state (15, 16).

In addition to regulating the RsmA network, NahK has also been implicated in NO-dependent processes in *P. aeruginosa*. *In vitro* studies have previously shown that NahK’s co-cistronic sensory protein NosP is a heme protein that can ligate picomolar concentrations of NO and lead to biofilm dispersal. However, this NO-mediated biofilm dispersal is unsuccessful in a *nahK* disrupted *P. aeruginosa* PAO1 strain (17, 18). More recent studies in a *P. aeruginosa* PA14 strain have also shown that NahK is necessary for sustainable denitrification, a process by which bacteria carry out respiration in an anaerobic environment (19, 20). When PA14 wildtype (WT) strain is grown in an oxygen depleted environment, denitrification reductases are upregulated during early stationary phase, to aid in energy generation by utilizing nitrate and nitrite as electron acceptors and converting them to nitrogen gas in a series of reductions. This leads to the accumulation of NO as an intermediate, which results in cell filamentation and favors anaerobic biofilm formation (19, 21). However, in a *nahK* deletion strain, this cell filamentation is observed much earlier in the growth phase, where there are presumably low levels of NO accumulated due to low cell density. Even during aerobic growth, slight cell filamentation is observed in the Δ*nahK* strain when exposed to nanomolar levels of exogenous NO, while no such filamentation is observed in the WT strain. This suggests that the Δ*nahK* strain may be more sensitive to signaling concentrations (pico- to nanomolar) of NO than the WT strain (19).

Given the multiple roles attributed to NahK in NO-dependent processes such as denitrification and NO-mediated biofilm dispersal, it is important to systematically understand the role of NosP/NahK in responding to NO (17, 19). Thus, in this work, the NO response of the Δ*nahK* strain is characterized and compared to that of WT, and the results observed indicate that NahK is required to protect *P. aeruginosa* from signaling concentrations of NO, which can otherwise lead to DNA damage induced SOS stress. This role of NahK is especially crucial in amino acid starved media and ultimately provides insight into previously unaddressed roles of NahK in *P. aeruginosa* metabolism.

## Results

### PA14 Δ*nahK* strain is susceptible to signaling concentrations of NO in an amino acid starved medium

The ubiquity of *P. aeruginosa* is attributed to its versatile metabolism and adaptability (22–24). This is enabled not only by the numerous two-component systems encoded in its genome, but also by the Gac MKN, which integrates multiple environmental signals to modulate RsmA-dependent phenotypes in this bacterium (14, 17, 25–27). Interestingly, phenotypes such as biofilm formation and virulence factor production have been shown to be influenced by essential nutrients such as iron, carbon, sulfur and phosphorus (28–30). The phenotypic differences of *P. aeruginosa* strains in different growth conditions necessitates the study of the bacterium to be conducted in a chemically defined medium to assess the role of medium composition in modulating the behavior of the bacterium. Hence, to determine the tolerance of PA14 WT and Δ*nahK* strains to NO, we grew both the strains in 50 mL of M9 minimal media supplemented with 0.4% (w/v) glucose as the carbon source and further subjected the cells to iron and nutrient deprivation. We then recorded the growth curves of the strains in both the presence and absence of ∼100 µM NO. In the absence of NO, both WT and Δ*nahK* strains displayed similar growth profiles. However, in the presence of NO, both strains displayed a longer lag phase, following which the NO-treated WT strain grew similar to the untreated WT strain, while the NO-treated Δ*nahK* strain showed a significant decrease in growth compared to the untreated Δ*nahK* strain (**Fig. 1A**). This was contrary to the previous results observed in rich media under aerobic conditions, and suggests that the sensitivity of Δ*nahK* strain to NO may be media dependent (19).

**Figure 1.**
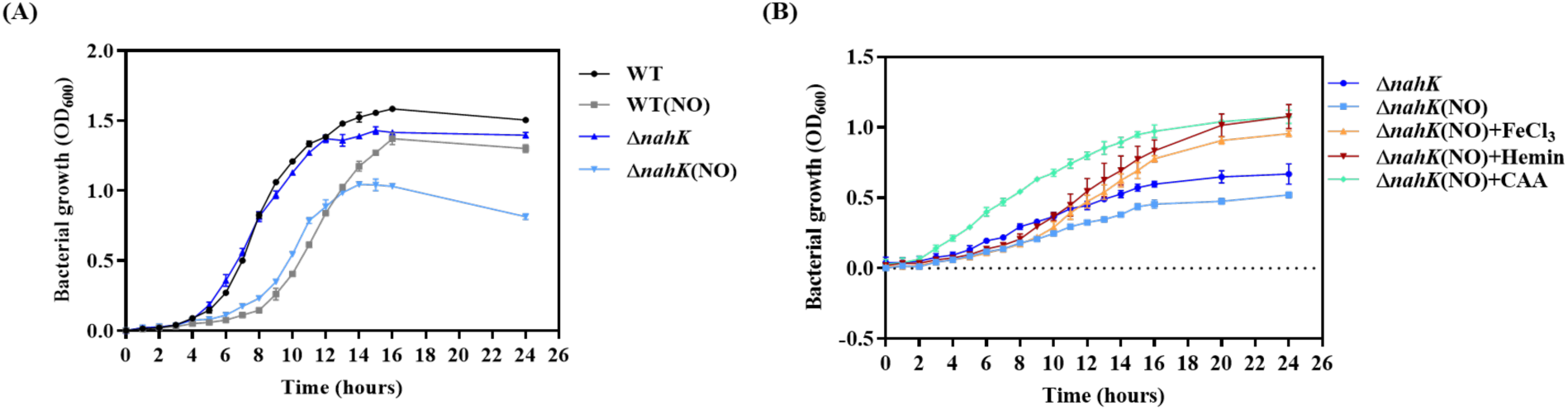
Δ*nahK* strain is sensitive to nanomolar concentrations of NO in a minimal medium. **(A)** Growth curves of PA14 WT and Δ*nahK* strains cultured in 50 mL of M9 minimal media supplemented with glucose as a carbon source, in the presence and absence of 100 μM DETA NONOate (which results in ∼100 nM NO over the course of growth). Error bars represent the standard deviation from the mean of values obtained from two independent experiments; some error bars may be smaller than the symbols. **(B)** Growth curves of Δ*nahK* strain in 5 mL of minimal media in the presence and absence of 50 μM DETA NONOate (which results in ∼50 nM NO over the course of growth), and Δ*nahK* strain treated with 50 μM DETA NONOate in 5 mL of minimal media supplemented with 5 μM FeCl_3_, 10 μM hemin, or 0.5% (w/v) casamino acids (CAA). Error bars represent the standard deviation from the mean of values obtained from three independent experiments; some error bars may be smaller than the symbols.

To evaluate how the components of the rich LB media may influence the susceptibility of Δ*nahK* strain to NO, we repeated the growth curve experiment in minimal media supplemented with nutrients in the form of amino acids and iron. Amino acids were added to the media in the form of 0.5% (w/v) casamino acids (CAA), and iron was added in the form of 5 μM FeCl_3_ or 10 μM hemin. While supplementation of iron significantly improved the growth of ΝΟ-treated Δ*nahK* strain during the exponential phase, the presence of CAA had a more robust effect with an early onset of exponential growth in just 2 hours (**Fig. 1B**). Thus, we hypothesized that upon deletion of *nahK* from PA14 WT and treatment with signaling concentrations of NO (pico-nanomolar levels), the bacteria may be experiencing a stress that is at least partially alleviated in the presence of a mixture of amino acids or iron sources in the media. Hence, we reasoned that NahK may be playing a role in alleviating nitrosative stress in *P. aeruginosa*.

### NO-stress causes a DNA damage-induced morphological change in the Δ*nahK* strain

NO is a signaling molecule at low concentrations (pico-nanomolar) but can have antimicrobial effects at high concentrations (millimolar) (31, 32). Higher concentrations of NO kill bacteria due to the generation of reactive nitrogen species (RNS), such as peroxynitrite, upon reacting with reactive oxygen species (ROS), such as hydroxyl radicals, which can damage the DNA, proteins, and lipids of bacteria (31, 33, 34). Lower concentrations of NO, such as the nanomolar levels endogenously generated through denitrification, is more tolerable for bacteria, albeit resulting in stress-induced phenotypes such as cell filamentation (21, 35). This phenotype may be a result of compromised DNA synthesis or NO-mediated DNA damage, leading to stalled replication and cell division (36, 37). To determine whether the Δ*nahK* strain is prone to NO-stress in minimal media, we imaged both WT and Δ*nahK* strains grown with and without NO using brightfield microscopy and quantified the average cell length of bacteria from each condition. Microscopy images revealed that WT strain retained normal cell morphology under both NO-treated and untreated conditions. However, the NO-treated Δ*nahK* strain had a significant population of filamented cells compared to the untreated Δ*nahK* strain, confirming the hypothesis that Δ*nahK* is subjected to nitrosative stress when cultured in minimal media in the presence of NO (**Fig. 2A and B**).

**Figure 2.**
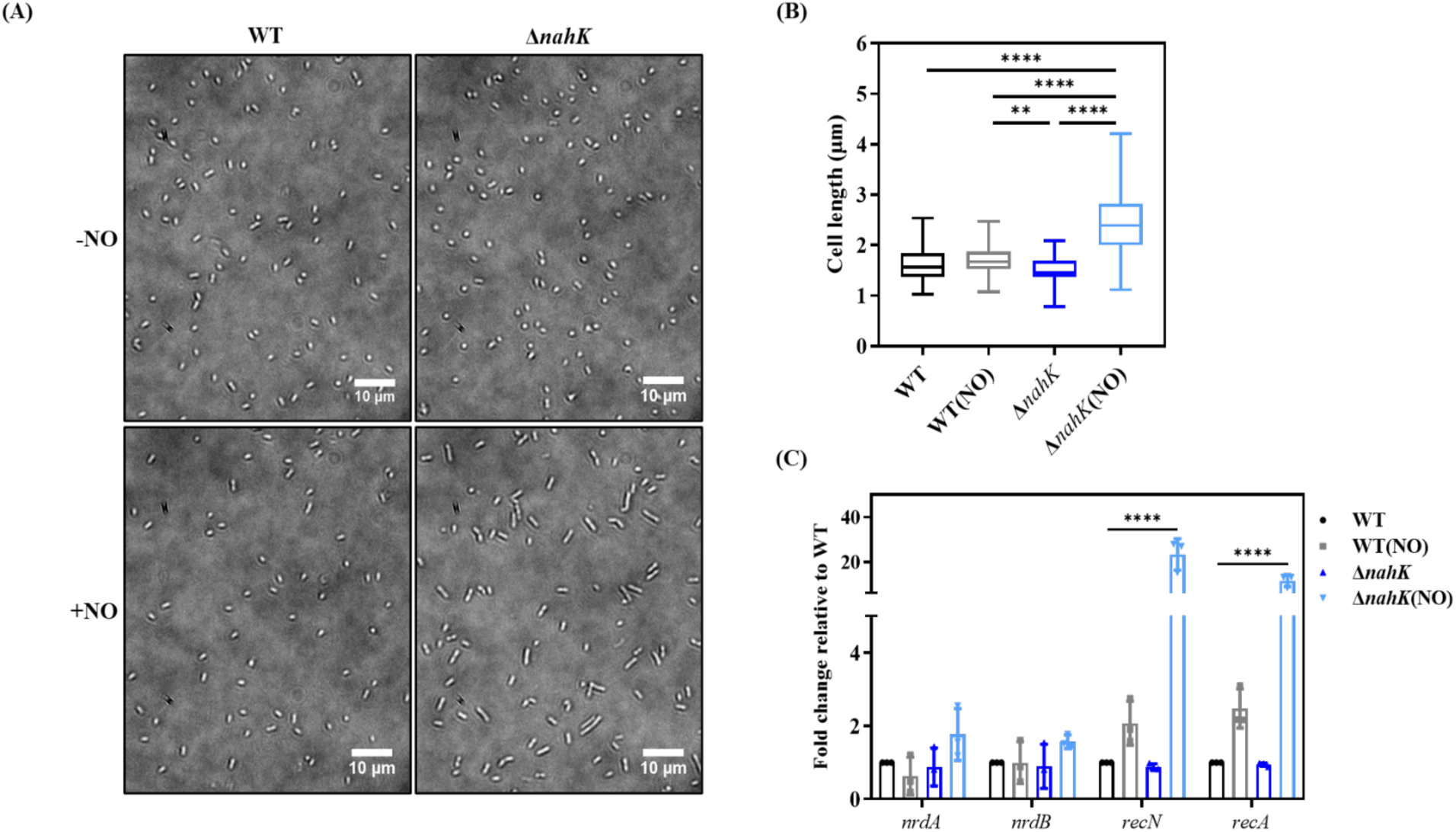
NO induced DNA damage causes cell filamentation in Δ*nahK* strain. **(A)** Representative microscopy images of WT and Δ*nahK* strains treated with or without 50 μM DETA NONOate, in 5 mL of minimal media. **(B)** Quantification of cell length of 30 randomly selected cells from 3 independent experiments each, using the line tool on ImageJ software. n=90. *p*-values were calculated using one-way analysis of variance (ANOVA) and Tukey’s multiple comparisons test. \*\**p* < 0.01 and \*\*\*\**p* < 0.0001. **(C)** qRT-PCR analysis of genes encoding the ribonucleotide reductases *nrdA* and *nrdB*, which are involved in DNA synthesis, and genes encoding the recombinases *recN* and *recA*, which are involved in DNA double strand break repair. *proC* (delta 1-pyrroline-5-carboxylate reductase) was used as the housekeeping gene and the fold change of each gene in all four conditions is reported relative to its expression in the untreated WT condition. Error bars represent the standard deviation from the mean of values obtained from three independent experiments, with two technical replicates each. Genes with fold changes greater than or less than 1 are considered upregulated or downregulated, respectively. *p*-values were calculated using two-way ANOVA and Dunnett’s multiple comparisons test comparing the mean ΔΔCt of a gene in all conditions to its mean ΔΔCt in the untreated WT culture. \*\*\*\**p* < 0.0001.

To determine whether the cell filamentation phenotype was caused by arrested cell division due to DNA damage, quantitative reverse transcription PCR (qPCR) was used to probe for the mRNA levels of genes involved in DNA damage repair in NO-treated and untreated WT and Δ*nahK* strains. For all qPCR analysis, total cellular RNA was extracted from large scale 50 mL cultures grown in the presence or absence of 100 μΜ DETA NONOate, to allow for sufficient growth of cells. To test for DNA damage, we determined the mRNA levels of *recN* and *recA*, as these genes encode for recombinase enzymes that repair DNA double strand breaks (38, 39). Both *recN* and *recA* were significantly upregulated in the NO-treated Δ*nahK* strain compared to the untreated Δ*nahK* strain, whereas no significant upregulation was observed in the NO-treated WT strain compared to untreated WT strain. To test whether cell filamentation was caused by insufficient DNA synthesis resulting in impaired cell division, we also tested the expression levels of *nrdA* and *nrdB*, which encode for *Pseudomonas* class I ribonucleotide reductases that are required for DNA synthesis under aerobic conditions (40). Neither *nrdA* nor *nrdB* showed any upregulation upon NO treatment in WT or Δ*nahK strains*. These genes were also not differentially regulated in untreated Δ*nahK* strain compared to untreated WT strain, indicating that DNA synthesis was not inherently compromised in the former (**Fig. 2C**).

As amino acids and iron sources enhanced the growth of NO-treated Δ*nahK* strain, we also tested the NO-stress induced cell filamentation phenotype under these conditions. Supplementation of iron to the growth media in the form of hemin and FeCl_3_ had no effect on cell phenotype whereas addition of amino acids in the form of CAA to the growth media completely complemented the filamented cell morphology in the NO-treated Δ*nahK* strain compared to that of untreated Δ*nahK* strain (**Fig. 3A and B**). This was correlated to lack of DNA damage, as neither *recN* nor *recA* were upregulated in the NO-treated Δ*nahK* strain cultured in media supplemented with CAA, suggesting that in amino acid starved media, the Δ*nahK* strain is more susceptible to NO mediated DNA damage (**Fig. 3C**).

**Figure 3.**
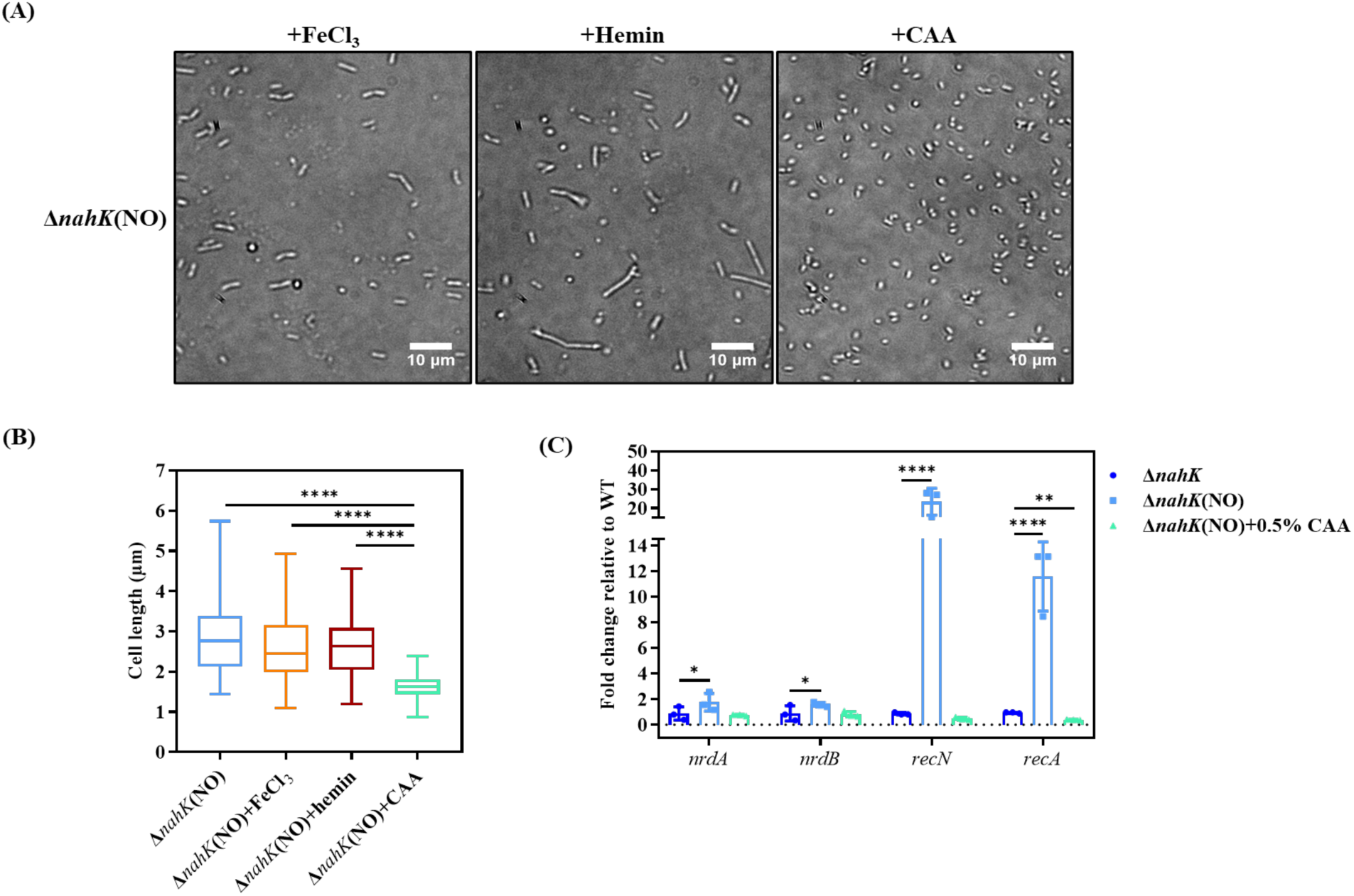
NO induced DNA damage and cell elongation in Δ*nahK* strain is alleviated in the presence of amino acids. **(A)** Representative microscopy images of the Δ*nahK* strain treated with 50 μM DETA NONOate, in minimal media supplemented with 5 μM FeCl_3_, or 10 μM hemin, or 0.5% (w/v) CAA. **(B)** Quantification of cell length of 30 randomly selected cells from 3 independent experiments each, using the line tool on ImageJ software. n=90. Cell lengths of NO-treated Δ*nahK* culture were also plotted for comparison (microscopy image not shown). *p*-values were calculated using one-way ANOVA and Tukey’s multiple comparisons test. \*\*\*\**p* < 0.0001. **(C)** qRT-PCR analysis of genes which encode for the ribonucleotide reductases *nrdA* and *nrdB*, and the recombinases *recN* and *recA*. *proC* was used as the housekeeping gene and the fold change of each gene in all three conditions is reported relative to its expression in the untreated WT condition, which is denoted by the dashed line at y=1. The relative fold changes of genes in the NO treated and untreated Δ*nahK* cultures were replotted from Fig.2 for comparison. Error bars represent the standard deviation from the mean of values obtained from three independent experiments, with two technical replicates each. Genes with fold changes greater than or less than 1 are considered upregulated or downregulated, respectively. *p*-values were calculated using two-way ANOVA and Dunnett’s multiple comparisons test comparing the mean ΔΔCt of a gene in all conditions to its mean ΔΔCt in the untreated Δ*nahK* culture. \**p* < 0.05, \*\**p* < 0.01 and \*\*\*\**p* < 0.0001.

### NO-stress causes cell lysis and eDNA release in the Δ*nahK* strain

When we harvested the WT and Δ*nahK* strains, cultured overnight with and without DETA NONOate, we observed that the NO-treated Δ*nahK* strain generated a cell pellet with a distinct slimy or sticky phenotype. Unlike the cell pellets generated from the untreated WT and Δ*nahK* strains or the NO-treated WT strain, this cell pellet could not be easily resuspended in the media (**Fig. 4A**). This slimy texture was reminiscent of the mucoid phenotype that is often associated with PA14 lung infections of cystic fibrosis patients (41–43). The mucoid phenotype is caused by the overproduction of a poly-uronic acid known as alginate, which is also one of the main virulence factors of *P. aeruginosa* and an essential component of its extracellular polymeric matrix (41, 44). Hence, we used qPCR to measure the expression levels of the *algD* gene, which is the first gene in the alginate biosynthesis operon. In the absence of NO, *algD* expression was similar in the WT and Δ*nahK* strains. However, in the presence of NO, *algD* was 30-fold upregulated in the Δ*nahK* strain while no significant upregulation was observed in the WT strain (**Fig. S1A**). We also attempted to determine the amount of alginate accumulated in each culture through a previously established uronic acid-carbazole assay, using an alginate standard (45). However, the experiment was unsuccessful in corroborating the results from the qPCR data due to the interference of glucose present in the minimal media (data not shown). Hence, the gene encoding AlgU, which is a sigma factor that facilitates the transcription of the alginate biosynthesis operon, was deleted from Δ*nahK* strain, to determine whether the sticky pellet phenotype was due to alginate overproduction (41). While deleting *algU* significantly attenuated the upregulation of *algD* in the NO-treated Δ*nahK*Δ*algU* strain compared to the NO-treated Δ*nahK* strain, it still generated a sticky cell pellet upon harvesting cells (**Fig. S1B**), suggesting that although alginate biosynthesis operon was upregulated and may have partially contributed to the slimy pellet phenotype, it was not the sole cause.

**Figure 4.**
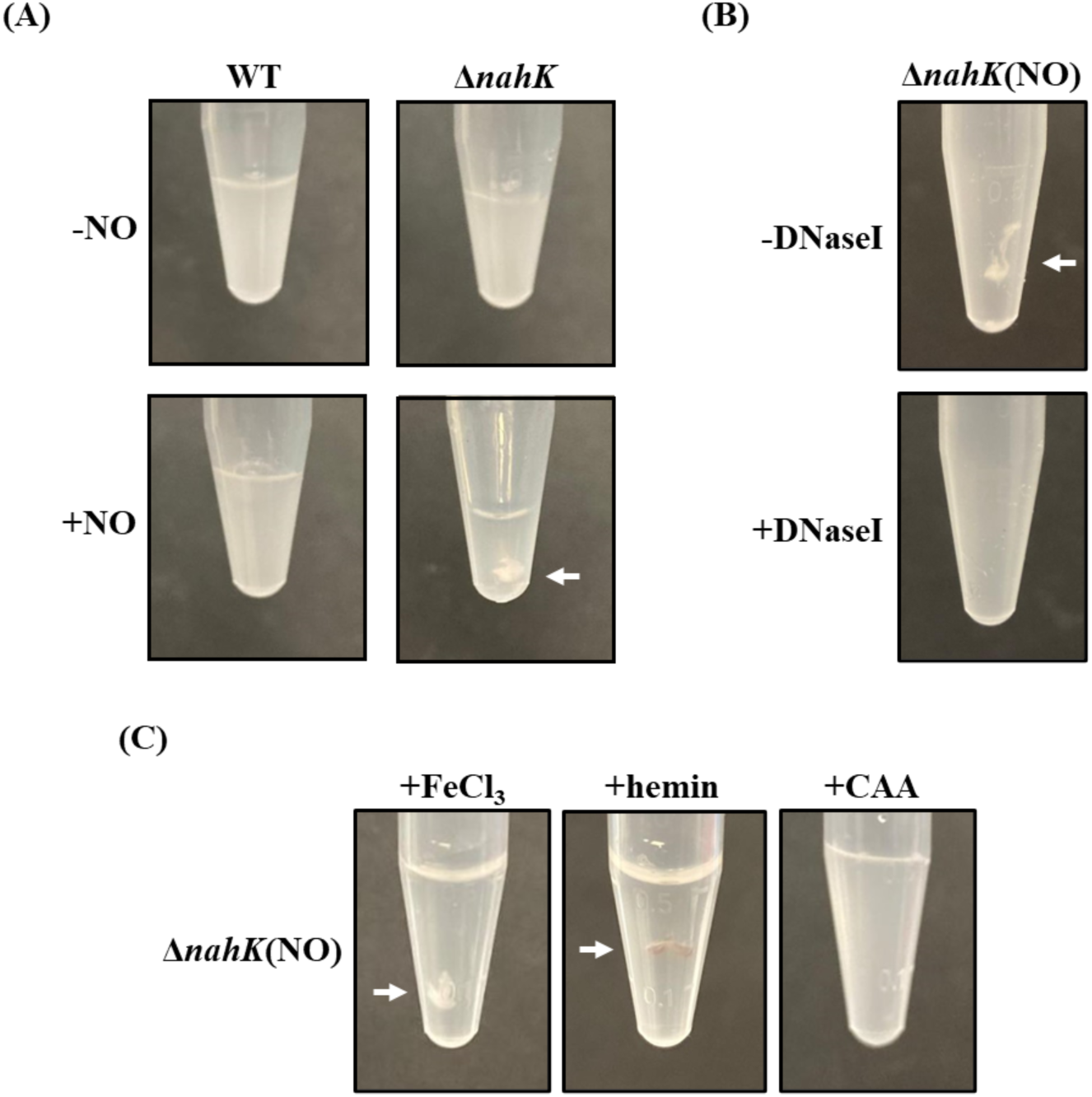
NO treated Δ*nahK* strain accumulates eDNA which results in the formation of slimy cell pellets. Representative images of resuspended pellets generated by harvesting cells from **(A)** 5 mL cultures of WT and Δ*nahK* strains grown in minimal media with or without 50 μM DETA NONOate, 5 mL cultures of Δ*nahK* strain treated with 50 μM DETA NONOate in minimal media supplemented **(B)** with or without 100 KU of DNaseI, or **(C)** with 5 μM FeCl_3_, or 10 μM hemin, or 0.5% (w/v) CAA. The images depict the formation of cell pellets which were either easily resuspended in media or were slimy in nature (indicated by white arrows) and could not be fully dispersed in media.

Another essential component of the biofilm matrix that contributes to its sticky texture is extracellular DNA (eDNA), which has been shown to be integral in maintaining biofilm architecture, through interactions with the exotoxin pyocyanin (46, 47). To check whether the sticky pellet phenotype was due to eDNA accumulation, the NO-treated Δ*nahK* strain was incubated with 100 KU of DNaseI for 15 min at room temperature prior to harvesting. Upon DNaseI incubation, the NO treated Δ*nahK* cell pellet was no longer sticky (**Fig. 4B**), suggesting that NO treatment caused cell lysis and the release of DNA in Δ*nahK* strain. To confirm that this cell lysis was also associated with NO mediated stress, the pellet texture of the NO-treated Δ*nahK* culture grown in minimal media supplemented with CAA was also evaluated, since amino acids can revert NO-stress induced cell filamentation in Δ*nahK* strain. Supplementation of CAA to culturing media completely prevented cell lysis in NO-treated Δ*nahK* culture, confirming that cell lysis was also caused by NO induced stress in the form of DNA damage. As expected, addition of FeCl_3_ or hemin had no effect on the cell pellet texture of NO-treated Δ*nahK strain*, just like in the case of cell filamentation (**Fig. 4C**).

We also quantified extracellular DNA accumulation in the NO-treated Δ*nahK* strain using propidium iodide (PI), which is a red fluorescent DNA intercalating stain that is not membrane permeable and hence can only stain the DNA in dead cells or DNA outside the cells, i.e., eDNA. To avoid PI fluorescence from staining dead cells, we performed a rapid attachment assay, wherein live bacteria were allowed to attach to chamber slides by incubating for a short period of time (15 minutes) and dead bacteria, which would be unable to attach due to lack of active pili, would be washed away. The attached bacteria in each chamber were then incubated with PI, followed by imaging. For each strain, we captured both red fluorescence microscopy images and corresponding brightfield microscopy images. The fluorescence microscopy images displayed several areas of red fluorescence in the NO-treated Δ*nahK* strain, indicating the presence of eDNA in this sample whereas little to no fluorescence was observed in the other samples (**Fig. 5A**). These areas were considered as eDNA accumulation sites if the corresponding areas in the respective brightfield images displayed cell clusters, as eDNA was found to cause cells to aggregate or clump together. Upon quantification of the number of eDNA sites in each culture, a significantly higher amount of eDNA accumulation was observed in the NO-treated Δ*nahK* culture, compared to all the other cultures (**Fig. 5C**). Upon supplementation of the NO-treated Δ*nahK* culture with DNaseI or upon culturing in the presence of CAA, the number of eDNA sites observed was notably decreased (**Fig. 5B and C**), substantiating the conclusion that NO-stress causes cell lysis in Δ*nahK* strain, which can be attenuated by the presence of amino acids in the culturing medium.

**Figure 5.**
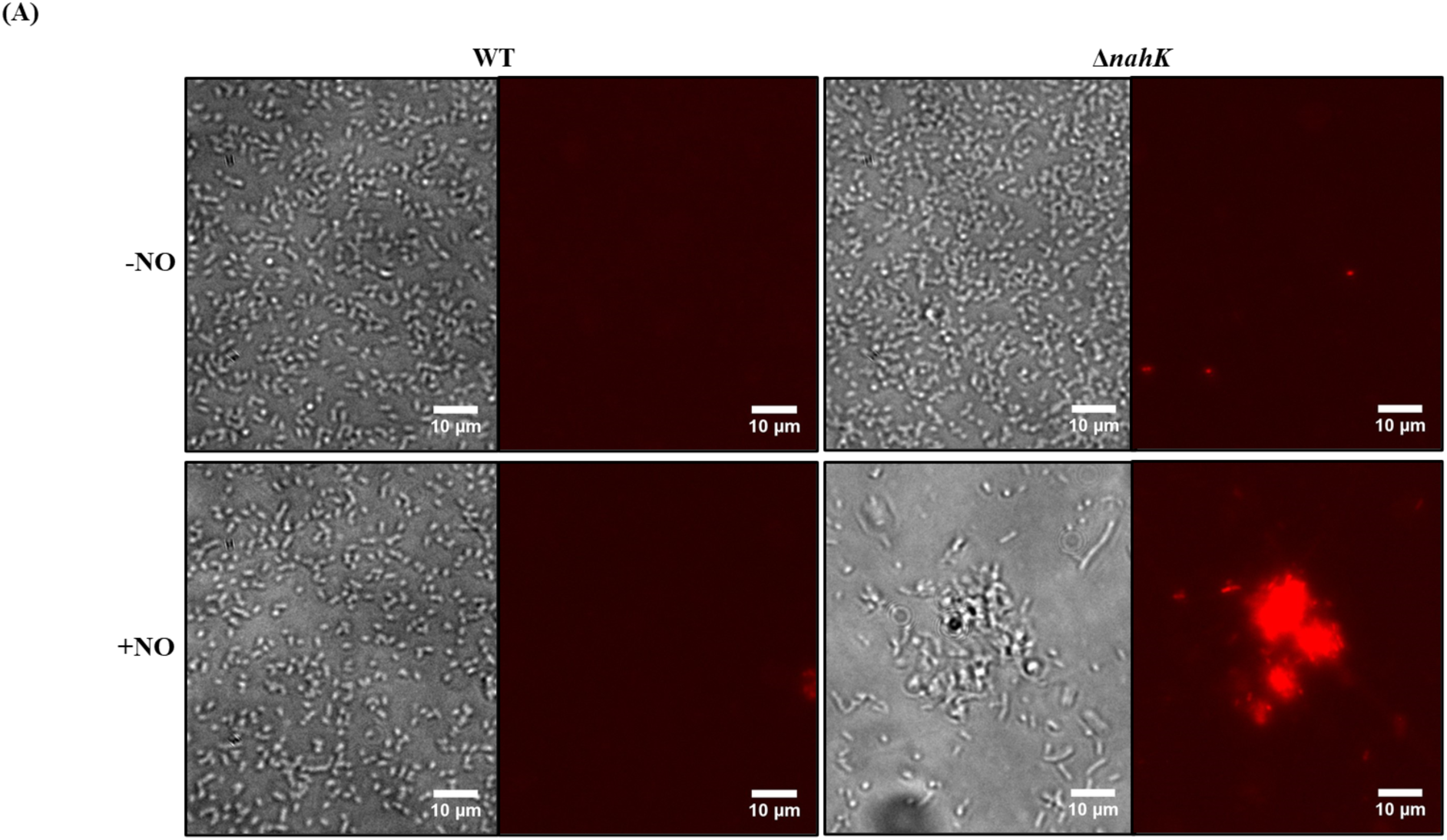
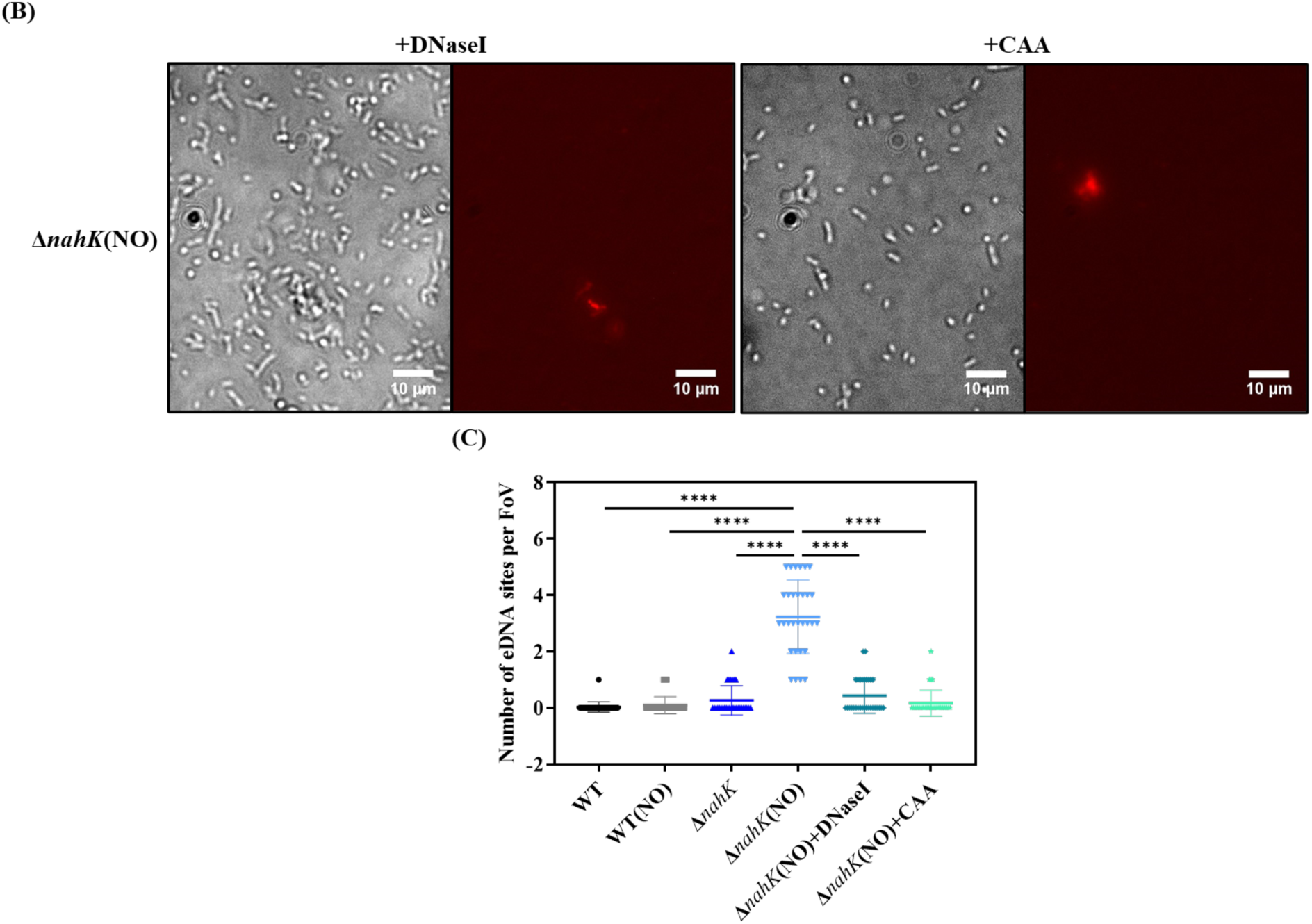
NO-stress induces eDNA release in Δ*nahK* strain. Representative brightfield (left) and corresponding fluorescence (right) microscopy images of propidium iodide (PI) stained **(A)** WT and Δ*nahK* strains cultured in 5 mL of minimal media in the presence or absence of 50 μM DETA NONOate and **(B)** NO-treated Δ*nahK* strain cultured in 5 mL minimal media supplemented with 100 KU DNaseI or 0.5% (w/v) CAA. A rapid attachment assay, where live cells were allowed to attach and dead cells were washed away, was used to obtain these images. The images show the presence or absence of PI-stained eDNA, indicated by red fluorescence in the fluorescence microscopy image, and the cell cluster formation facilitated by eDNA in the corresponding brightfield microscopy image. **(C)** Number of eDNA sites observed per field of view (FoV) in a fluorescence microscopy image, that corresponds to cell clusters in the respective brightfield microscopy image. The values were determined from a total of 30 random FoVs (n=30) obtained from 3 independent experiments. *p*-values were calculated using one-way ANOVA and Tukey’s multiple comparisons test. \*\*\*\**p* < 0.0001.

### NO treatment induces the SOS stress response in the Δ*nahK* strain

To understand the differences in the response of both WT and Δ*nahK* strains to NO, we conducted total RNA sequencing of both the strains, cultured with and without NO. From the results obtained, genes were considered to be significantly upregulated only if their log_2_ fold change was greater than or equal to one, with a *p*-value less than 0.05. Similarly, genes were considered to be significantly downregulated only if their log_2_ fold change was less than or equal to one, with a *p*-value less than 0.05. Out of the 6022 genes in the PA14 genome, only 91 genes were differentially expressed in NO-treated WT strain compared to the untreated WT strain, out of which 69 were significantly upregulated and 22 were significantly downregulated (**Fig. 6B**, **Tables S4, S5, S7 and S8**). Whereas a total of 1337 genes were differentially expressed in NO-treated Δ*nahK* strain compared to the untreated Δ*nahK* strain, out of which 629 were significantly upregulated and 708 were significantly downregulated (**Fig. 6C, Tables S4, S6, S7 and S9**). Interestingly, out of all the differentially expressed genes, only 33 genes were commonly upregulated and 7 genes were commonly downregulated, in both NO-treated WT and NO-treated Δ*nahK* strains compared to their untreated counterparts (**Tables S4 and S7**).

**Figure 6.**
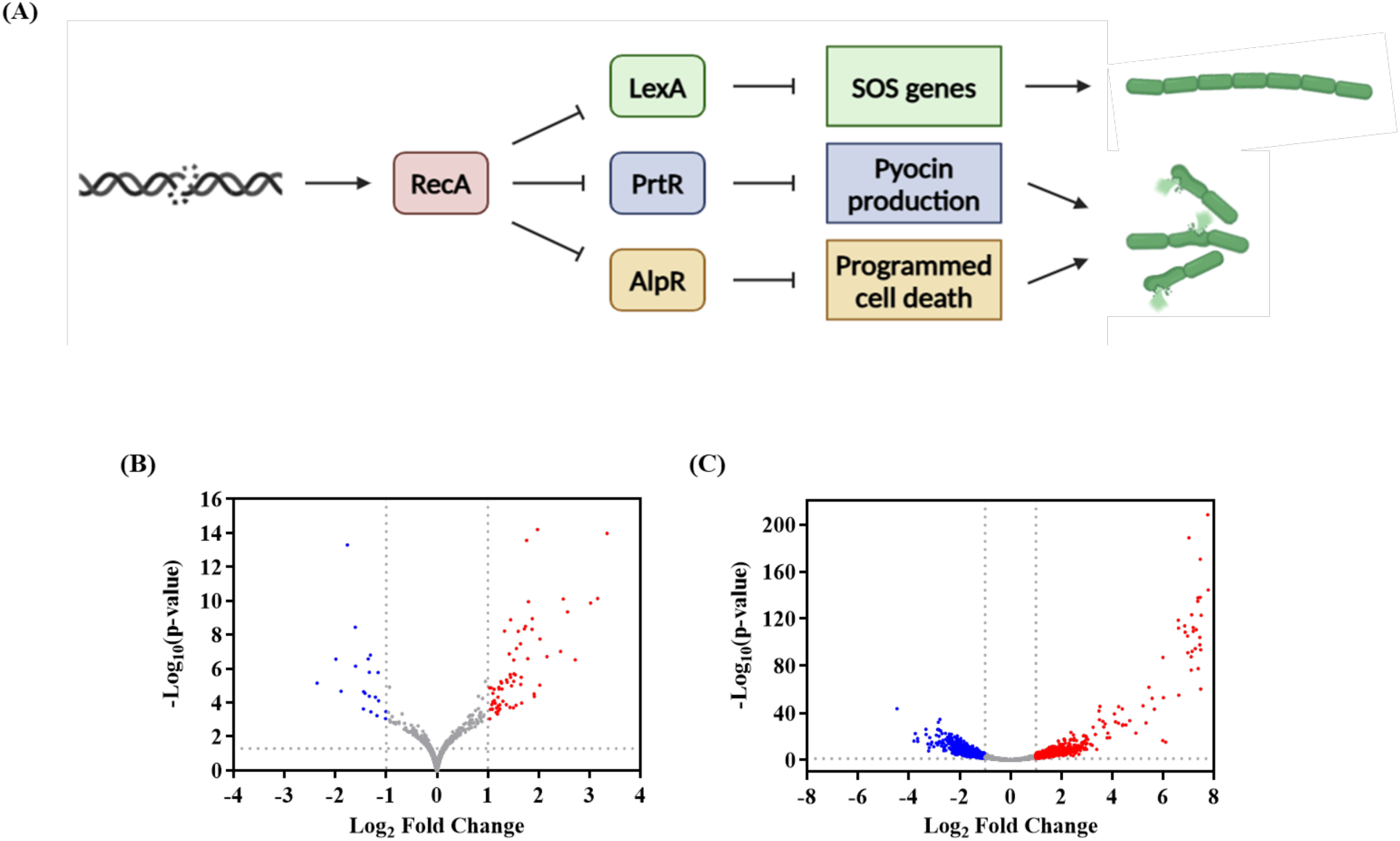
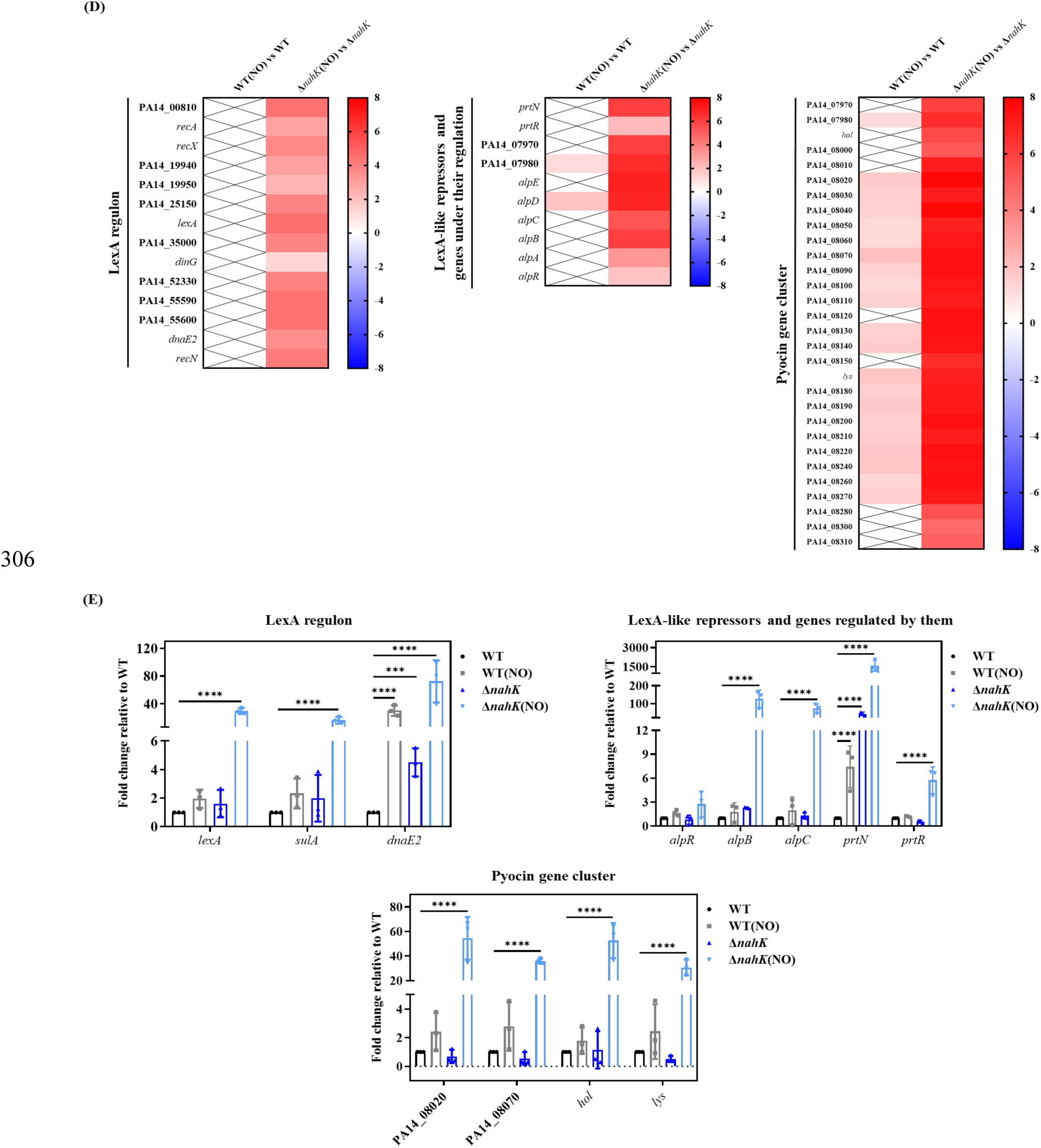
SOS stress response is activated in Δ*nahK* strain upon exposure to nanomolar concentrations of NO. (A) Schematic representation of the SOS stress response pathway in P. aeruginosa. Volcano plot depicting differentially regulated genes in **(B)** NO treated WT strain vs untreated WT strain, and **(C)** NO-treated Δ*nahK* strain vs untreated Δ*nahK* strain. Red data points indicate genes that were significantly upregulated with a log_2_ fold change greater than 1 and −log_10_(*p*-value) greater than 1.301 (*p*-value < 0.05). Blue data points indicate genes that were significantly downregulated with a log_2_ fold change less than −1 and −log_10_(*p*-value) greater than 1.301 (*p*-value < 0.05). All other data points, that were not significant, are depicted in gray. **(D)** Heat maps displaying the log_2_ fold changes of canonical SOS stress response genes and pyocin genes differentially regulated in NO treated vs untreated WT and Δ*nahK* strains, with *p*-value < 0.05. **(E)** qRT-PCR analysis of select SOS stress response genes in WT and Δ*nahK* strains grown in 50 mL minimal media in the presence or absence of 100 μM DETA NONOate. *proC* was used as the housekeeping gene and the fold change of each gene in all three conditions is reported relative to its expression in the untreated WT condition. Error bars represent the standard deviation from the mean of values obtained from three independent experiments, with two technical replicates each. Genes with fold changes greater than or less than 1 are considered upregulated or downregulated, respectively. *p*-values were calculated using two-way ANOVA and Dunnett’s multiple comparisons test comparing the mean ΔΔCt of a gene in all conditions to its mean ΔΔCt in the untreated WT culture. \*\*\**p* < 0.001 and \*\*\*\**p* < 0.0001.

Out of the significantly upregulated genes in the NO-treated Δ*nahK* strain compared to the untreated Δ*nahK* strain, the most upregulated genes belong to the pyocin cluster of PA14 (**Fig. 6D, Table S6**). Pyocins or bacteriocins are antimicrobials produced and released by *P. aeruginosa* to fight closely related species in an environment where both strains may need to compete for resources (48). Pyocin production is often associated with the bacterial SOS stress response, which is triggered by damage to DNA and involves the initiation of expression of proteins involved in DNA damage repair (49–51). Interestingly, cell filamentation and cell lysis of bacteria are also phenotypes that are often associated with the bacterial SOS stress response (**Fig. 6A**) (50, 52). Hence, it was hypothesized that NO-treatment of Δ*nahK* strain might be inducing the SOS stress response due to the DNA damage caused by the RNS that may be generated in this strain upon exposure to nanomolar levels of NO. Analysis of the RNA sequencing results confirmed this hypothesis, as canonical SOS stress response genes were significantly upregulated in the NO-treated Δ*nahK* strain compared to the untreated Δ*nahK* strain, whereas none of these genes were differentially expressed in the NO-treated WT strain compared to the untreated WT strain (**Fig. 6D, Tables S5 and S6**) (53).

These genes are part of the regulon of a repressor LexA, which undergoes autocleavage upon activation of the recombinase RecA, which in turn is caused by the binding of RecA with single stranded DNA fragments generated from DNA double strand breaks (**Fig. 6A**) (53). As stated earlier, some of these genes, such as *recA, recN, recX, dinG* and *dnaE2,* are involved in repairing this DNA damage, in an error free and error prone manner, resulting in advantageous mutations for the bacterium. The LexA regulon also includes *sulA* (PA14_25150), which encodes for the protein that inhibits cell division and results in cell filamentation during SOS stress response. Upregulated along with the LexA regulon, are genes encoding LexA-like repressors AlpR (PA14_52530) and PrtR (PA14_07960), which negatively regulate the expression of genes involved in programmed cell death (PCD) and pyocin production respectively (**Fig. 6A**) (53–55). It is possible that, like LexA, these repressors are also cleaved due to the NO triggered SOS stress in Δ*nahK* because genes involved in PCD (PA14_52520 or *alpA*, PA14_52510 or *alpB*, PA14_52500 or *alpC*, PA14_52490 or *alpD* and PA14_52480 or *alpE*) and pyocin production (PA14 07950 or *prtN* and PA14_07970 to PA14_08310) were also upregulated in NO-treated Δ*nahK* compared to untreated Δ*nahK* culture (**Fig. 6D, Table S6**). 32 out of the 36 genes in the *P. aeruginosa* pyocin gene cluster were more than 8-fold upregulated in these conditions, and this also comprised the genes encoding holin Hol (PA14_07990) and endolysin Lys (PA14_08160), which have been previously implicated in explosive cell lysis in interstitial biofilms, to facilitate pyocin release (56). Similarly, *alpB* is an annotated holin that also facilitates cell lysis during PCD by forming pores in the cell membrane (55, 57). Thus, the eDNA release observed in the NO-treated Δ*nahK* strain must be due to cell lysis caused by PCD or pyocin release. Interestingly, some of the genes in the pyocin gene cluster were also significantly upregulated in the NO-treated WT strain compared to the untreated WT strain, although to a lesser extent, indicating that there may be some DNA damage caused by NO in the WT strain as well (**Fig. 6D, Table S4**). To verify these results, we used qPCR to determine the expression levels of select SOS stress response genes belonging to three categories; genes belonging to the LexA regulon, genes that encode for LexA-like repressors and are regulated by them, and genes that are part of the pyocin gene cluster. qPCR data corroborated the results from RNA sequencing and confirmed that SOS stress response was triggered in NO-treated Δ*nahK* culture (**Fig. 6E**).

To confirm whether the cell filamentation and lysis phenotypes observed in the NO-treated Δ*nahK* strain were due to DNA damage induced SOS stress response, we treated both WT and Δ*nahK* strains with ciprofloxacin (CPX), a known SOS inducing drug that causes DNA damage by targeting DNA gyrase (53). Both CPX-treated WT and Δ*nahK* strains mimicked the cell filamentation and lysis phenotypes of the NO-treated Δ*nahK* strain (**Fig. 7**). Interestingly, unlike NO-stress, CPX-stress could not be alleviated by CAA, also suggesting that amino acids may be preventing NO-stress by neutralizing the source of stress, that is RNS, and not by directly repairing DNA damage (**Fig. S2**). Taken together, our data confirms that NO-treatment of Δ*nahK* strain causes DNA damage and triggers the SOS stress response, resulting in cell filamentation and cell lysis.

**Figure 7.**
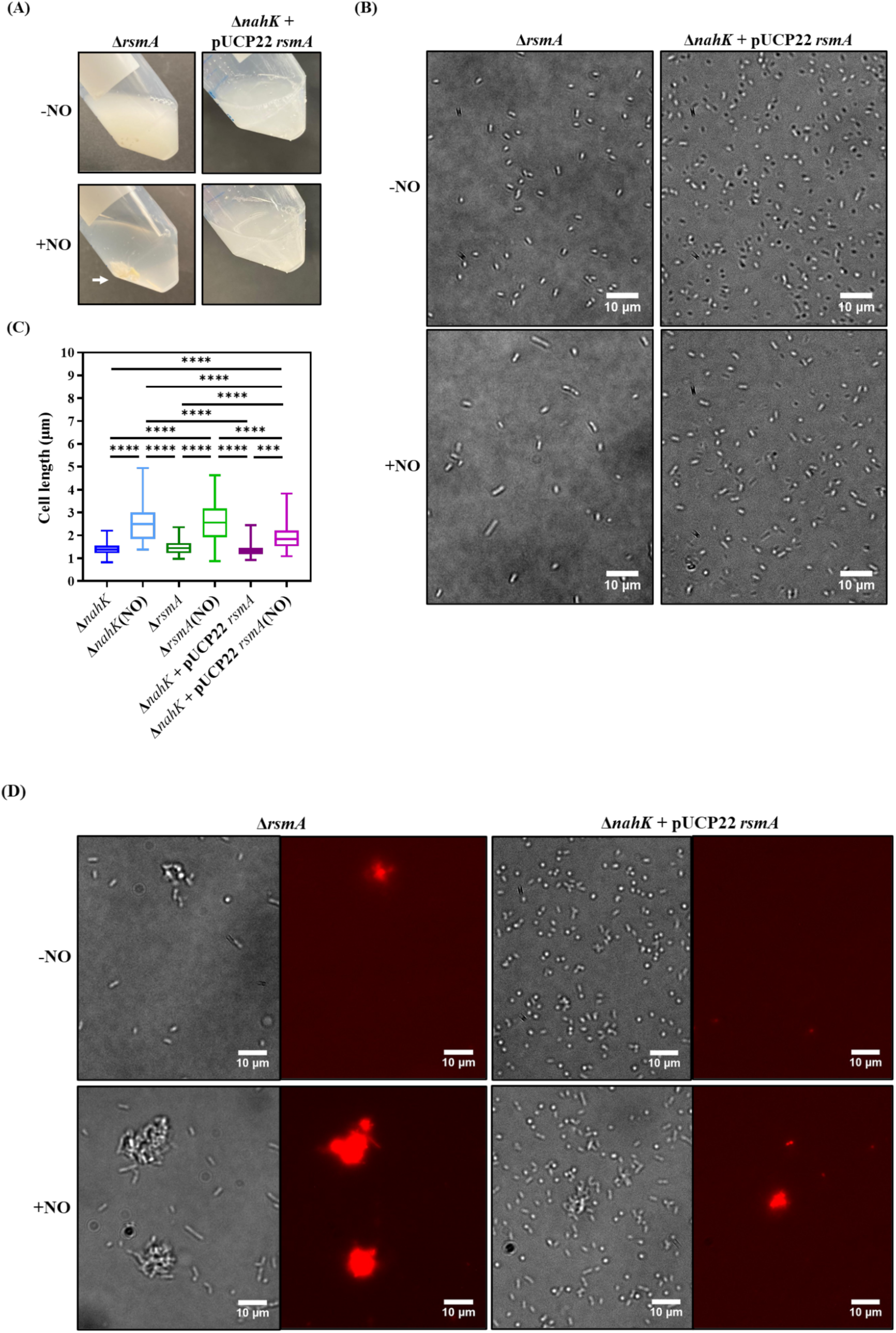
Cell filamentation and lysis phenotypes observed in NO-treated Δ*nahK* strain is due to activation of the SOS stress response. **(A)** Representative images of resuspended pellets generated by harvesting cells from 50 mL cultures of WT and Δ*nahK* strains treated with 0.025 μg/mL ciprofloxacin (CPX), a known SOS inducing drug, in minimal media. The images depict the formation of thick slimy cell pellets (indicated by white arrows) that could not be fully resuspended in media. **(B)** Representative microscopy images of WT and Δ*nahK* strains treated with 0.025 μg/mL CPX in 5 mL of minimal media. **(C)** Quantification of cell length of 30 randomly selected cells from 3 independent experiments each, using the line tool on ImageJ software. n=90. Cell lengths of untreated WT and Δ*nahK* cultures were also plotted for comparison (microscopy images not shown). *p*-values were calculated using one-way ANOVA and Tukey’s multiple comparisons test. \*\*\*\**p* < 0.0001.

### RsmA activity can protect the Δ*nahK* strain from NO-stress

NahK has been shown to be an important regulator of the switch between biofilm and planktonic lifestyles in *P. aeruginosa* PA14 owing to its effect on the activity of the global post-transcriptional regulator protein RsmA. Deletion of the *nahK* gene from PA14 WT stimulates the overproduction of the exotoxin pyocyanin, attenuates its motility, and results in the formation of more robust biofilms, all of which are phenotypes associated with a chronic infection lifestyle of *P. aeruginosa* (15, 16). Deletion of *nahK* also mis-regulates denitrification under anaerobic conditions due to the decreased transcription of denitrification reductases (19). All these phenotypes have been associated with the inhibition of RsmA activity; the Δ*rsmA* strain phenotypically mimics the Δ*nahK* strain and overexpression of *rsmA* in Δ*nahK* complements all these phenotypes. Therefore, it is important to determine if the susceptibility of the Δ*nahK* strain to NO stress is also due to an inhibition of RsmA activity in this strain. Hence, we grew both the Δ*rsmA* and the *rsmA* overexpressed Δ*nahK* strains (Δ*nahK*+pUCP22 *rsmA*) in M9 minimal media with glucose as the carbon source, supplemented with or without DETA NONOate. After overnight growth, the NO-treated Δ*rsmA* strain displayed all the SOS stress phenotypes associated with the NO-treated Δ*nahK* strain and overexpression of *rsmA* in Δ*nahK* partially reverted all these phenotypes to that of WT. NO-treatment of the Δ*rsmA* strain induced DNA damage, as validated by the formation of filamented cells due to inhibited cell division and by the upregulation of *recN* and *recA* by 40-fold and 15-fold, respectively. Interestingly, the ribonucleotide reductase genes *nrdA* and *nrdB* were also significantly upregulated, further confirming DNA damage in the NO-treated Δ*rsmA* strain. However, in the *rsmA* overexpressed Δ*nahK* strain, fewer filamented cells were observed upon NO treatment and *recN* and *recA* were only half as upregulated as in the NO-treated Δ*nahK* culture (**Fig. 8B, C and F**). Upon harvesting the cells, the NO-treated Δ*rsmA* culture generated a sticky pellet similar to that of the NO-treated Δ*nahK* strain, whereas the *rsmA* overexpressed Δ*nahK* strain generated a cell pellet which was neither sticky like the NO-treated Δ*nahK* cell pellet, nor easily dissolvable in media like the NO-treated WT cell pellet (**Fig. 8A**). Similarly, NO treatment of the Δ*rsmA* strain resulted in more eDNA accumulation sites whereas overexpression of *rsmA* in Δ*nahK* significantly attenuated eDNA release in this strain (**Fig. 8D and E**). Δ*nahK* carrying an empty overexpression plasmid (pUCP22 empty) still retained its cell filamentation and lysis phenotypes upon NO treatment, confirming that the partial complementation of phenotypes in *rsmA* overexpressed Δ*nahK* was solely due to increased expression of RsmA that is presumably in its active or ON state (**Fig. S3**).

**Figure 8.**
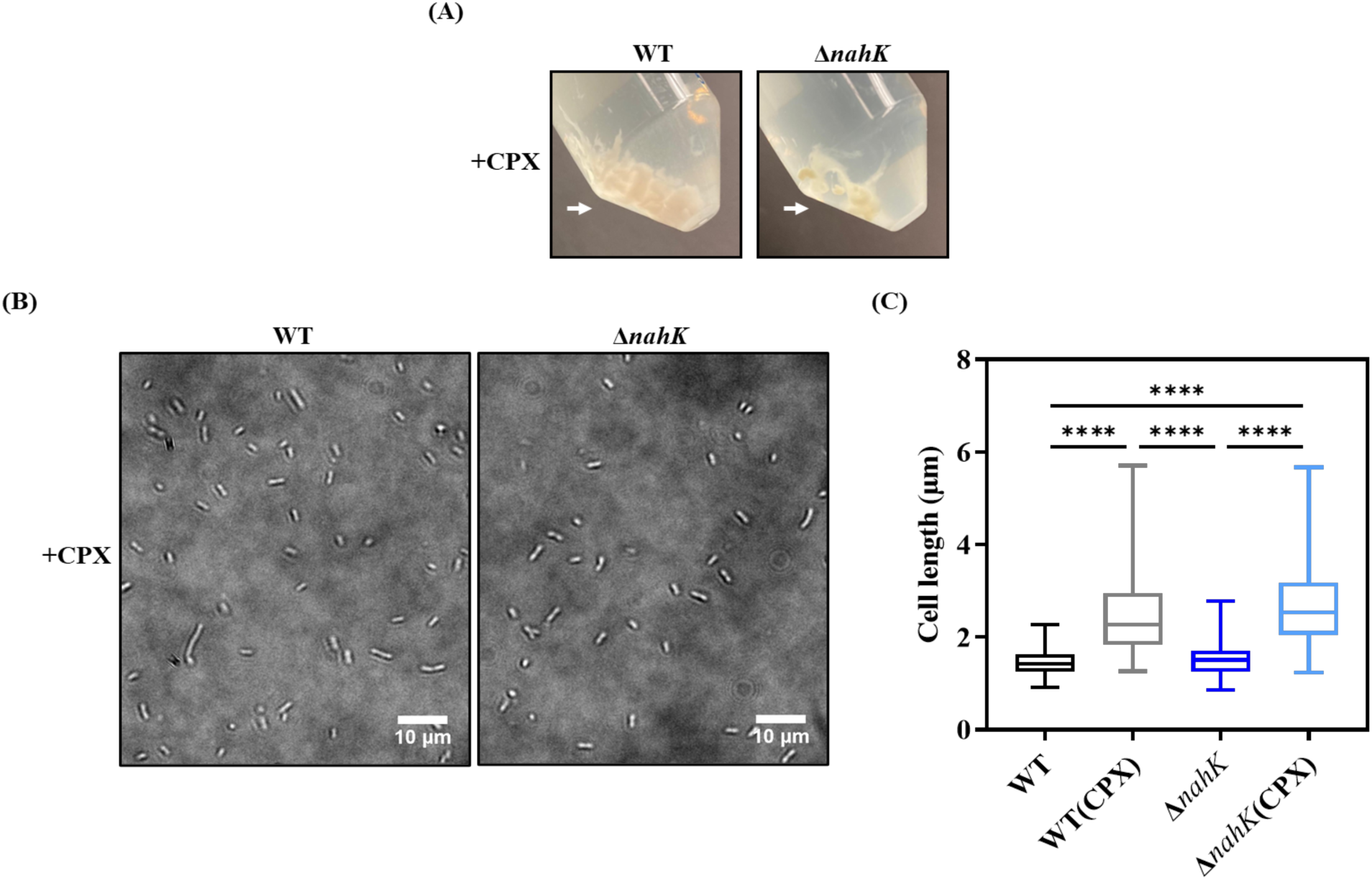
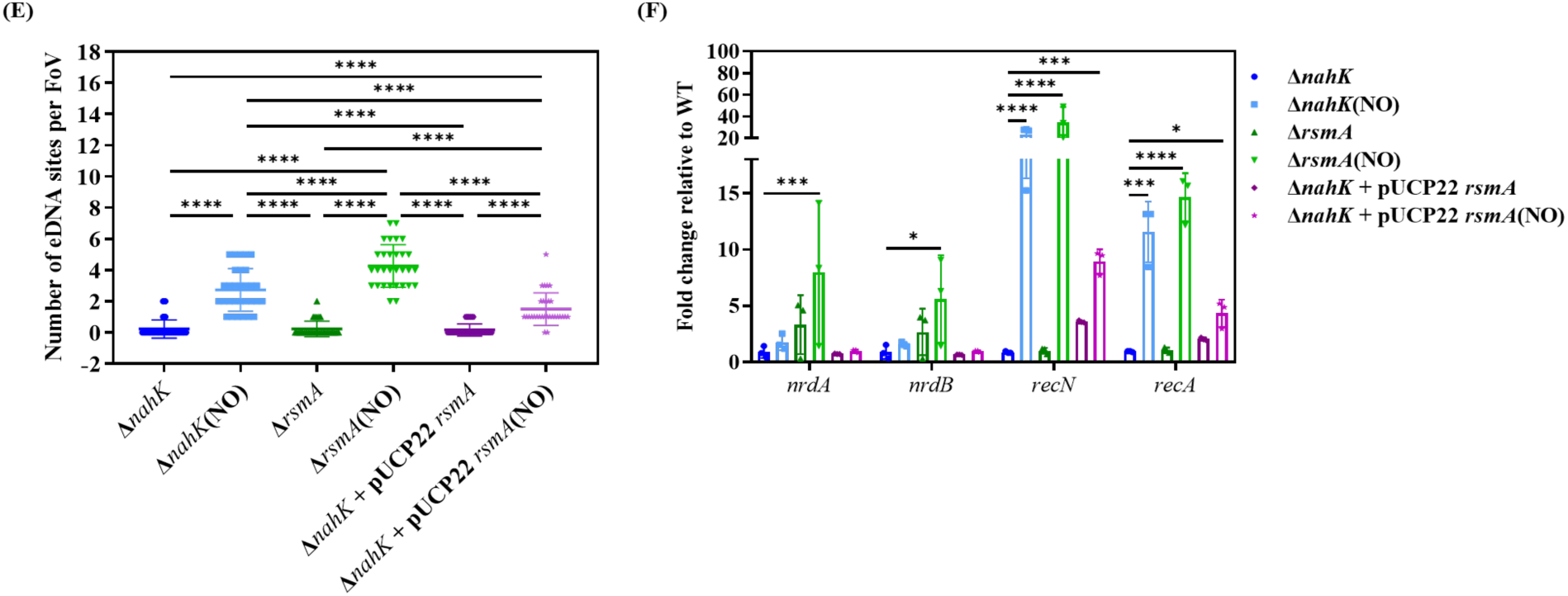
Hypersensitivity of Δ*nahK* strain to NO might be due to inhibition of RsmA. **(A)** Representative images of resuspended pellets generated by harvesting cells from 50 mL cultures of both the Δ*rsmA* and the *rsmA* overexpressed Δ*nahK* strains, cultured in minimal media in the presence or absence of 100 μM DETA NONOate. The images depict the formation of dissoluble cell pellets upon harvesting both the untreated strains, and the formation of a thick and slimy cell pellet in the case of the NO treated Δ*rsmA* strain (indicated by a white arrow). The cell pellet generated from the NO treatment of the *rsmA* overexpressed Δ*nahK* strain had an intermediary texture that was not discernible in the images captured. **(B)** Representative microscopy images of the Δ*rsmA* and *rsmA* overexpressed Δ*nahK* strains cultured with or without 50 μM DETA NONOate, in 5 mL of minimal media. **(C)** Quantification of cell length of 30 randomly selected cells from 3 independent experiments each, using the line tool on ImageJ software. n=90. Cell lengths of NO-treated and untreated Δ*nahK* cultures were also plotted for comparison (microscopy images not shown). *p*-values were calculated using one-way ANOVA and Tukey’s multiple comparisons test. \*\*\**p* < 0.001 and \*\*\*\**p* < 0.0001. **(D)** Representative brightfield (left) and corresponding fluorescence (right) microscopy images of propidium iodide stained Δ*rsmA* and *rsmA* overexpressed Δ*nahK* strains cultured in 5 mL minimal media in the presence or absence of 50 μM DETA NONOate. The images were generated using the rapid attachment assay and show the presence or absence of PI-stained eDNA (red fluorescence) in the fluorescence microscopy images, and cell cluster formation facilitated by eDNA in the corresponding brightfield microscopy images. **(E)** Number of eDNA sites observed per FoV in fluorescence microscopy images, that corresponded to cell clusters in the respective brightfield microscopy image. The values are generated from a total of 30 random FoVs (n=30) obtained from 3 independent experiments. The number of eDNA sites per FoV in the NO treated and untreated Δ*nahK* cultures were replotted from Fig.5 for comparison. *p*-values were calculated using one-way ANOVA and Tukey’s multiple comparisons test. \*\*\*\**p* < 0.0001. **(F)** qRT-PCR analysis of genes encoding ribonucleotide reductases and recombinases in NO treated and untreated Δ*nahK*, Δ*rsmA* and *rsmA* overexpressed Δ*nahK* strains. The relative fold changes of genes in the NO treated and untreated Δ*nahK* cultures were replotted from Fig.2 for comparison. *proC* was used as the housekeeping gene and the fold change of each gene in all six conditions is reported relative to its expression in the untreated WT condition, which is denoted by the dashed line at y=1. Error bars represent the standard deviation from the mean of values obtained from three independent experiments, with two technical replicates each. Genes with fold changes greater than or less than 1 are considered upregulated or downregulated, respectively. *p*-values were calculated using two-way ANOVA and Dunnett’s multiple comparisons test comparing the mean ΔΔCt of a gene in all conditions to its mean ΔΔCt in the untreated Δ*nahK* culture. \**p* < 0.05, \*\*\**p* < 0.001 and \*\*\*\**p* < 0.0001.

While we have established the relevance of the kinase function of NahK in regulating RsmA, NahK also possesses an annotated sensory domain called the Per-Arnt-Sim (PAS) domain, whose function is currently unknown (17). Given the role of amino acids in inhibiting NO-mediated stress in the Δ*nahK* strain, we also examined whether this sensory domain of NahK plays a role in its response to NO, through detection of amino acids. Hence, we generated a ΔPAS *nahK* strain by deleting the entire PAS domain of NahK, encoded by the first 459 amino acids. Upon assessing the mutant strain’s NO-responsive phenotypes, we observed that the cells were neither filamented nor underwent lysis (**Fig. S4**). This confirmed that the susceptibility of Δ*nahK* to signaling concentrations of NO in minimal media is exclusively due to its kinase activity resulting in RsmA being turned OFF and not due to any potential sensing function of its PAS domain.

### Increased susceptibility of the Δ*nahK* strain to NO may be due to sulfate starvation

From all the results observed thus far, we can conclude that the Δ*nahK* strain is susceptible to NO induced DNA damage, resulting in cell lysis and cell filamentation. It is very intriguing that as low as 50 nM NO, which is a concentration normally involved in signaling, can cause SOS stress in the Δ*nahK* strain. We considered several hypotheses to explain this. First, we considered that the Δ*nahK* strain may be already under redox imbalance or oxidative stress that is exacerbated by the addition of NO due to the generation of RNS. The most likely explanation for this would be the overproduction of pyocyanin in Δ*nahK*, as pyocyanin production can cause ROS accumulation and lead to cell lysis (16, 30). However, culturing the Δ*nahK* strain with NO in minimal media supplemented with M64, an inhibitor that has been previously shown to completely inhibit pyocyanin production, had no effect on cell filamentation or lysis (58). Surprisingly, overnight growth of Δ*nahK* with NO and an ROS scavenger Tiron also did not prevent NO-stress (**Fig. S5**).

We also considered the hypothesis that Δ*nahK* may be generating and accumulating NO due to misregulated denitrification, as shown in previous studies (19). To test this hypothesis, we incubated Δ*nahK* strain with the NO scavenger PTIO prior to overnight growth with DETA NONOate, to remove any NO that may have accumulated through denitrification. We also incubated Δ*nahK* overnight with NaNO_3_, a denitrification substrate, in order to test for endogenous NO generation through misregulated denitrification in Δ*nahK*. However, pre-incubation with PTIO did not inhibit NO-stress in Δ*nahK* and overnight incubation with NaNO_3_ alone did not stimulate NO-stress (**Fig. S6**).

We then considered the hypothesis that the Δ*nahK* strain is inefficient in detoxifying even the small amount of NO that it is exposed to in these experiments. This possibility was less likely, as the concentration of NO in the medium is not considered toxic, and the gene encoding the aerobic NO detoxification enzyme, Fhp, in *P. aeruginosa* is upregulated approximately 3-fold in both NO-treated WT and NO-treated Δ*nahK* cultures, compared to the untreated cultures, according to our RNA sequencing results (**Table S4**) (59). Nonetheless, to test this hypothesis, a *lacZ* reporter was generated for the *fhp* promoter (P*fhp*-*lacZ*) and inserted into both the WT and Δ*nosP*Δ*nahK* strains, the latter of which phenotypically mimics Δ*nahK* (**Fig. S7**) (insertion of plasmid into the Δ*nahK* was unsuccessful), and their β-galatosidase activities were evaluated. While the reporter in WT showed a small increase in activity upon NO-treatment, it was not statistically significant, and no such increase was observed for the reporter in the NO-treated Δ*nosP*Δ*nahK* strain compared to the untreated strain. We also tested the NO-stress phenotypes of an *fhp* deletion strain by growing Δ*fhp* in minimal media supplemented with or without DETA NONOate. However, deletion of *fhp* from PA14 had no effect on its tolerance to NO, as NO treatment of Δ*fhp* strain neither generated a sticky cell pellet nor resulted in formation of filamented cells (**Fig. S8**).

Despite these setbacks in understanding the NO-induced stress in Δ*nahK*, we persisted in finding an explanation. It is evident that increased susceptibility of the Δ*nahK* strain to NO can be completely reverted upon exposure to amino acids in the culturing media and partially alleviated by overexpressing *rsmA* in Δ*nahK* (**Fig. 3**, **Fig. 4**, **Fig. 5 and Fig. 8**). RsmA, being a global post-transcriptional regulator in *P. aeruginosa*, can modulate the translation of more than 500 downstream genes (13). Amino acids could either be promoting the de-repression of RsmA by influencing the transcription of *rsmY*/*Z*, or facilitating the translation of proteins that are essential in responding to low concentrations of NO. To test these possibilities, we cultured Δ*rsmA* strain in minimal media in the presence and absence of NO and CAA. NO-treated Δ*rsmA* culture neither produced filamented cells nor formed a slimy cell pellet upon harvesting, confirming that CAA could alleviate NO-stress in the Δ*rsmA* strain similar to the Δ*nahK* strain (**Fig. 9**). This suggests that CAA alleviates NO-stress by affecting the translation of proteins that are downstream of RsmA and not by de-repressing RsmA to promote its ON state.

**Figure 9.**
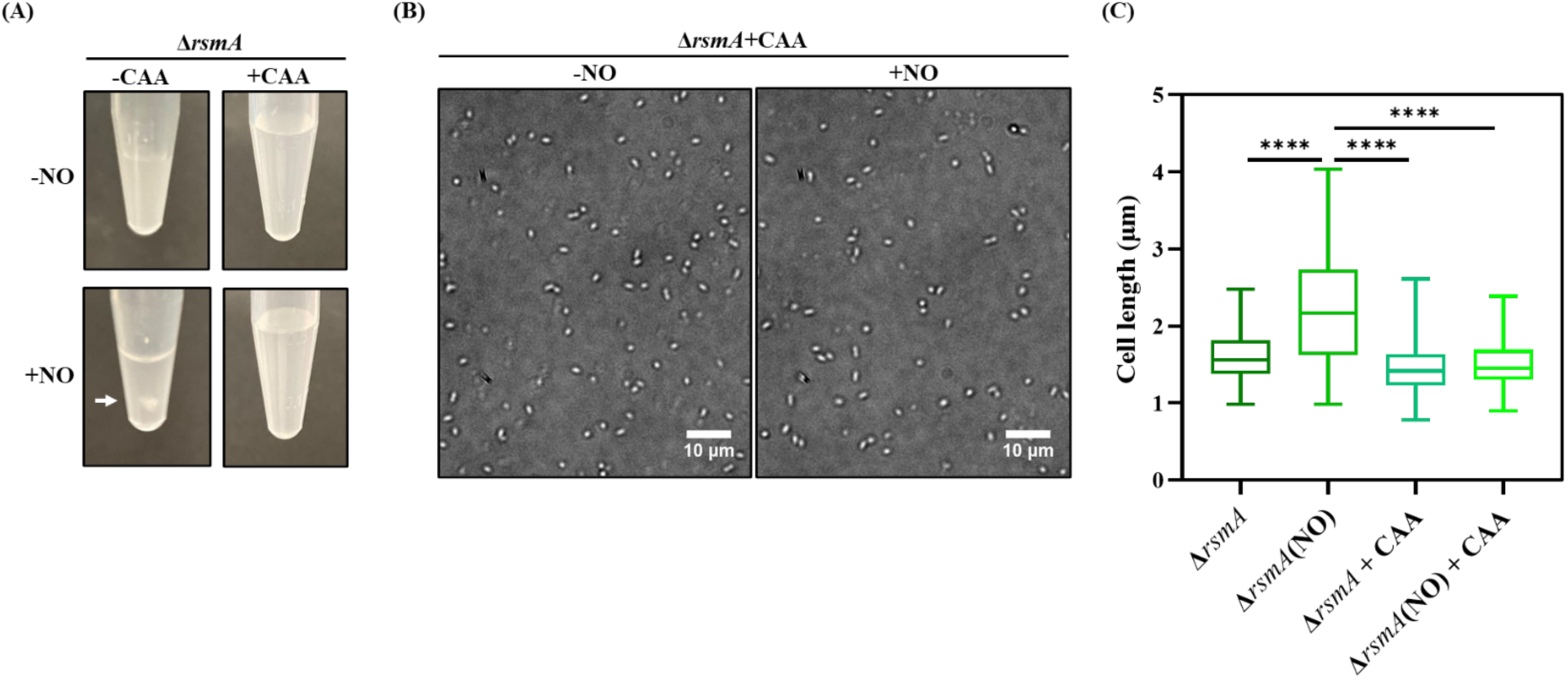
Supplementation of amino acids in the media can alleviate NO-stress in Δ*rsmA* strain. **(A)** Representative images of resuspended pellets generated by harvesting cells from 5 mL cultures of Δ*rsmA* strain treated with or without 50 μM DETA NONOate in minimal media supplemented with or without 0.5% (w/v) CAA. The images depict the formation of a slimy cell pellet (indicated by white arrow) in the NO treated Δ*rsmA* strain grown in amino acid deficient media and dispersible cell pellets in all the other conditions tested. **(B)** Representative microscopy images of Δ*rsmA* strain treated with or without 50 μM DETA NONOate in 5 mL of minimal media supplemented with 0.5% (w/v) CAA. **(C)** Quantification of cell length of 30 randomly selected cells from 3 independent experiments each, using the line tool on ImageJ software. n=90. Cell lengths of NO-treated and untreated Δ*rsmA* cultures were also plotted for comparison (microscopy images not shown). *p*-values were calculated using one-way ANOVA and Tukey’s multiple comparisons test. \*\*\*\**p* < 0.0001.

Previous transcriptomic studies of the Δ*rsmA* strain revealed that 706 genes were upregulated and 256 genes were downregulated in the Δ*rsmA* strain, relative to WT, when cultured in M9 minimal media supplemented with glucose as the carbon source (15). The genes that are upregulated or downregulated in Δ*rsmA*, relative to WT, are the genes that are negatively regulated or positively regulated, respectively, by RsmA. Negative regulation of RsmA entails binding of the protein to GGA motifs on the hexaloops of its target mRNAs, whereas positive regulation of RsmA does not involve a direct interaction of RsmA with target mRNAs, and may be achieved indirectly by the repression of a repressor (13). From **Fig. 9** it is evident that CAA protects Δ*rsmA* from NO-stress despite the protein not being expressed. Hence, it is possible that the genes involved in resisting NO stress may be the genes that are positively regulated by RsmA. This narrows down the total list of differentially regulated genes to the genes that are commonly downregulated in Δ*nahK* and Δ*rsmA* relative to WT. There are 153 genes that are commonly downregulated in both the strains with a log2 fold change of less than −1 and p-value of less than 0.05. These were functionally categorized according to their assigned pathways in Kegg (https://www.genome.jp/kegg/pathway.html) (**Fig. 10A**).

**Figure 10.**
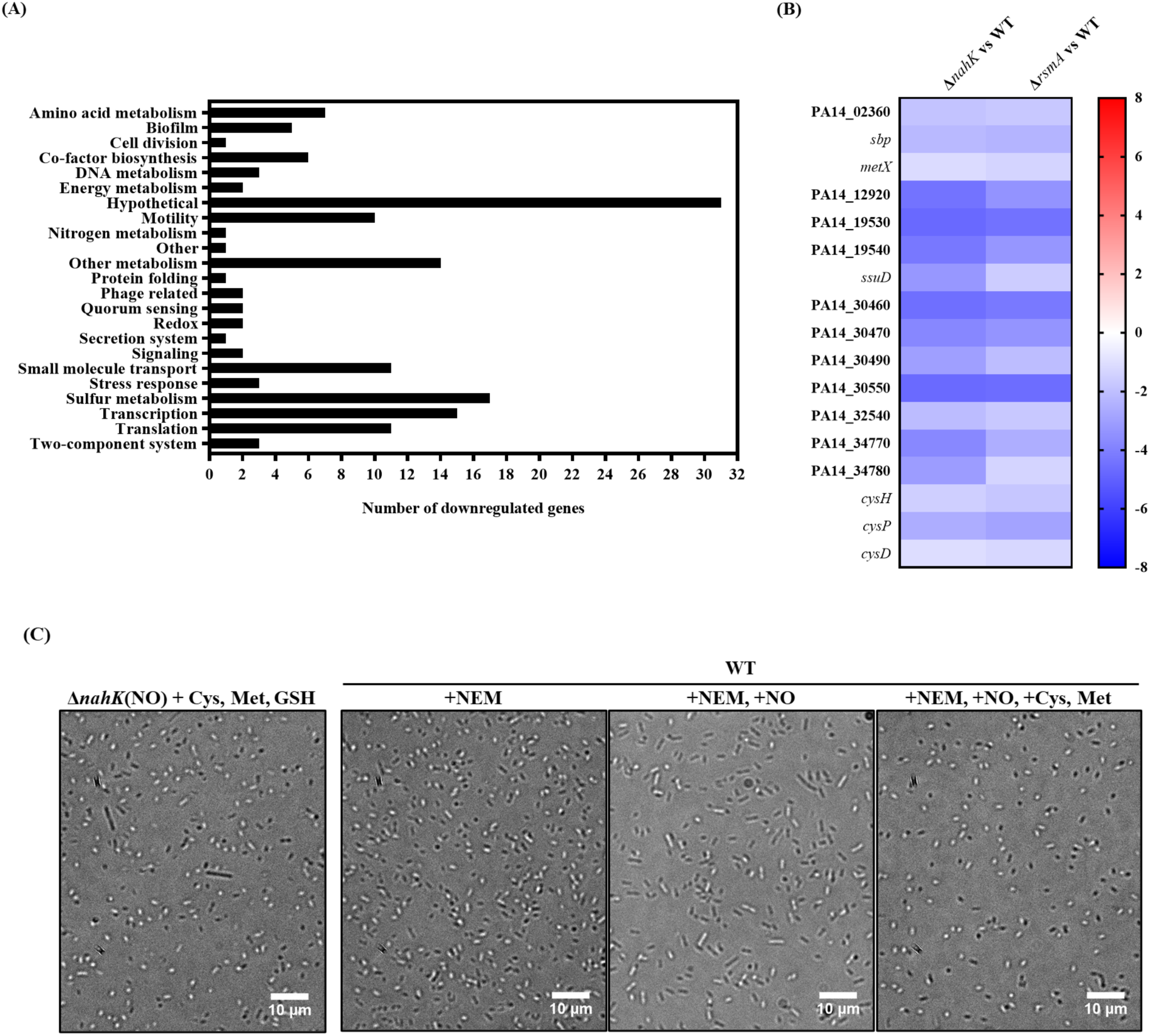
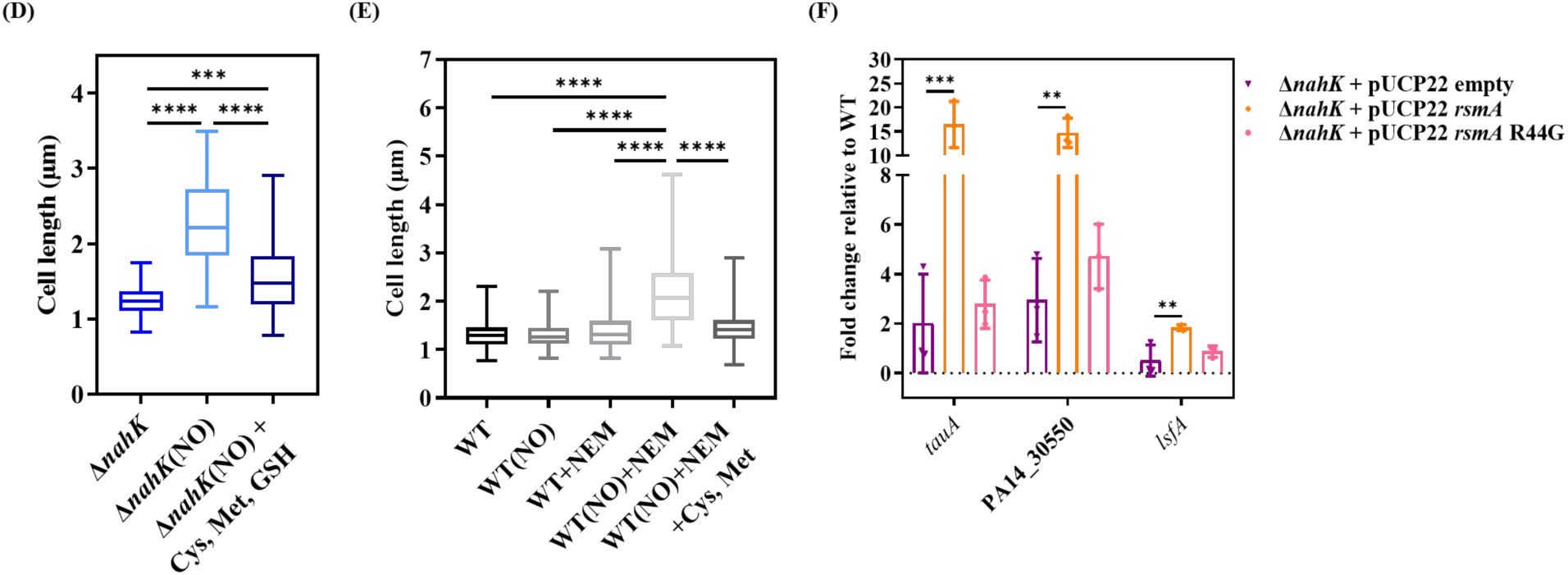
Susceptibility of Δ*nahK* to low concentrations of NO may be due to sulfate starvation. **(A)** Functional classification of genes which were more than one-fold downregulated in both the Δ*nahK* and the Δ*rsmA* strains relative to WT, with *p*-value less than 0.05, as reported in recent studies from our lab (15). **(B)** Heat maps displaying the log_2_ fold changes of sulfur metabolism genes downregulated in both Δ*nahK* and Δ*rsmA* strains relative to WT, with log_2_ fold change < −1 and *p*-value < 0.05. **(C)** Representative microscopy images of the untreated Δ*nahK* strain and the NO treated Δ*nahK* strain in the presence or absence of 1 mM Cys, Met and GSH, along with representative microscopy images of the WT strain treated with 50 μM NEM in the presence or absence of 50 μM DETA NONOate, in 5 mL of minimal media supplemented with or without 1 mM Cys and Met. **(D)** and **(E)** Quantification of cell length of 30 randomly selected cells from 3 independent experiments each, using the line tool on ImageJ software. n=90. Cell lengths of NO-treated and untreated WT and Δ*nahK* cultures are also plotted for comparison (microscopy images not shown). *p*-values were calculated using one-way ANOVA and Tukey’s multiple comparisons test. \*\**p* < 0.01 and \*\*\*\**p* < 0.0001. **(F)** qRT-PCR analysis of sulfate starvation induced genes in Δ*nahK* strains harbouring an empty pUCP22 plasmid, a plasmid overexpressing *rsmA* or a plasmid overexpressing an inactive mutant of *rsmA* (R44G). *proC* was used as the housekeeping gene and the fold change of each gene in all three strains is normalized to its expression in the WT strain. Error bars represent the standard deviation from the mean of values obtained from three independent experiments, with two technical replicates each. Genes with fold changes greater than or less than 1 are considered upregulated or downregulated, respectively. *p*-values were calculated using two-way ANOVA and Dunnett’s multiple comparisons test comparing the mean ΔΔCt of a gene in all conditions to its mean ΔΔCt in the Δ*nahK* strain harbouring empty pUCP22 plasmid. \*\**p* < 0.01 and \*\*\**p* < 0.001.

Interestingly, 17 genes involved in sulfur metabolism were commonly downregulated in both the strains relative to WT. More than half of these genes are involved in transport of sulfur sources such as sulfate/thiosulfate (*sbp*, *cysP*), sulfonate (PA14_02360, PA14_19540, PA14_30470, PA14_30550, PA14_34770, PA14_34780) and taurine (PA14_12920), while the remaining are involved in metabolism of sulfur sources (**Fig. 10B**) (60, 61). Sulfur is important for maintaining cellular redox, as it is an essential part of Fe-S cluster containing antioxidant enzymes. Compromised sulfur metabolism also affects cysteine (Cys) and methionine (Met) metabolism as sulfur is essential for the biosynthesis of both the amino acids, and this was indicated by the down regulation of genes involved in Cys and Met metabolism as well (*serA*, *metE*, *metH*) (62). Cysteine is also involved in alleviating cellular oxidative stress as it is a major component of glutathione (GSH), the most powerful cellular antioxidant, and is also required for the proper functioning of antioxidant enzymes such as thioredoxins, peroxiredoxins and glutaredoxins. These findings suggest that increased sensitivity of Δ*nahK* to NO-stress may be due to compromised sulfur metabolism leading to depreciated cellular thiol content.

To test this hypothesis, we cultured the Δ*nahK* strain with 50 μM NO in 5 mL minimal media containing a combination of 1 mM Cys, Met and GSH. After overnight incubation, cells were imaged under the microscope, followed by centrifugation to check the cell pellet texture. The pellet texture was neither as slimy as NO-treated Δ*nahK* nor as easily dissoluble like the untreated Δ*nahK*. Since a visibly sticky cell pellet was not generated, the data is not shown. However, upon imaging the cells under the light microscope, it became evident that the presence of sulfur containing amino acids was sufficient to almost fully alleviate NO-stress in Δ*nahK*, as almost all the cells possessed normal cell length and the population of even slightly filamented cells was lower than in the NO-treated Δ*nahK* strain (**Fig. 10C and D**). However, this effect was not specific to the presence of sulfur containing amino acids alone, as previously shown in **Fig. 3**, where a mixture of amino acids in the form of casamino acids could complement the filamentation phenotype in NO-treated Δ*nahK* strain. Similar results were also observed upon addition of different mixtures of other amino acids such as alanine (Ala), valine (Val), arginine (Arg) and glutamine (Gln) (**Fig. S9**). Interestingly, a higher concentration of a single amino acid such as GSH did not elicit similar effects, despite its antioxidant properties, suggesting that in the presence of a mixture of amino acids, SOS stress may be alleviated due to activation of other cellular defenses against NO stress.

To further confirm the role of amino acids in alleviating NO-stress, we treated the WT strain, with or without 50 μM NO, in 5 mL minimal media in the presence of 50 μM serine hydroxamate (SHX), to induce the stringent response by causing amino acid starvation (63). To specifically test the role of sulfur containing amino acids in reducing NO induced stress, the WT strain was also treated with or without 50 μM NO in the presence of 50 μM N-ethyl maleimide (NEM), which depletes cellular thiols by reacting with free thiol groups (64). We then tested the filamentation and cell clustering phenotypes by imaging and cell harvesting, respectively. The pellet texture of the harvested cells did not have any perceptible stickiness, unlike the NO-treated Δ*nahK* cultures. Hence, cell length quantification was used as a readout for NO-stress. In the presence of SHX or NEM alone, a small population of WT cells seemed to be longer than cells under non-stressed conditions. This effect was more pronounced upon treatment of the WT strain with NO in the presence of SHX, where a larger population of filamented cells was observed (**Fig. S10**). However, cell filamentation was most evident when Δ*nahK* was exposed to NO in the presence of NEM, with a greater population of visibly longer cells. To confirm that this increased susceptibility to NO in the presence of NEM was indeed due to thiol depletion, the experiment was repeated in minimal media supplemented with 1 mM Cys and Met. This reverted the cells back to normal cell length, confirming that in the absence of cellular thiols, even WT cells, which are otherwise resistant to nanomolar concentrations of NO, now possess increased sensitivity to NO, similar to the Δ*nahK* strain (**Fig. 10C and E**). We further tested whether RsmA positively regulates sulfur metabolism genes since overexpression of *rsmA* in the Δ*nahK* strain partially alleviated NO-stress in this strain. When compared to either the Δ*nahK* strain harboring an empty pUCP22 plasmid, or the Δ*nahK* strain overexpressing an inactive mutant of *rsmA* (R44G) which does not alleviate NO stress (**Fig. S11**), sulfur metabolism genes encoding transporters of sulfur sources such as taurine (*tauA*) and sulfonate (PA14_30550), as well as a thiol dependent antioxidant enzyme *lsfA,* were upregulated in the *rsmA* overexpressed Δ*nahK* strain (**Fig. 10F**) (15, 60, 61). These results suggest that the Δ*nahK* strain may be more susceptible to NO-stress due to its inability to combat oxidative/nitrosative stress due to compromised sulfur metabolism in this strain.

## Discussion

In this work, we highlight a previously unexplored role of the histidine kinase NahK in NO mediated stress resistance in *P. aeruginosa* (**Fig. 11**). A PA14 strain that lacks *nahK* is hypersensitive to nanomolar concentrations of NO in minimal media; such low concentrations are usually only involved in signaling, not stress. This hypersensitivity results in NO-mediated DNA damage, which activates the SOS stress response pathway, as shown by the upregulation of the LexA regulon. This response manifests in the form of phenotypes such as cell filamentation and lysis-mediated eDNA release. We also show that hypersensitivity of the Δ*nahK* strain to nanomolar concentrations of NO can be alleviated by either the overexpression of *rsmA* or by supplementation of amino acids to the media. While *rsmA* overexpression partially relieved NO-stress, amino acids completely prevented stress, suggesting that RsmA might be indirectly involved in positively regulating the translation of these amino acids. This was further supported by the downregulation of sulfur metabolism genes in the Δ*nahK* and Δ*rsmA* strains, relative to WT. Supplementation of sulfur containing amino acids such as Cys, Met and GSH partially protected Δ*nahK* strain from NO-stress, while depletion of cellular thiols from WT strain induced stress upon exposure to nanomolar concentration of NO. Thus NahK, which is necessary for NO signaling in *P. aeruginosa*, is also indirectly involved in resistance to stress induced by signaling concentrations of NO, through the regulation of RsmA.

**Fig. 11.**
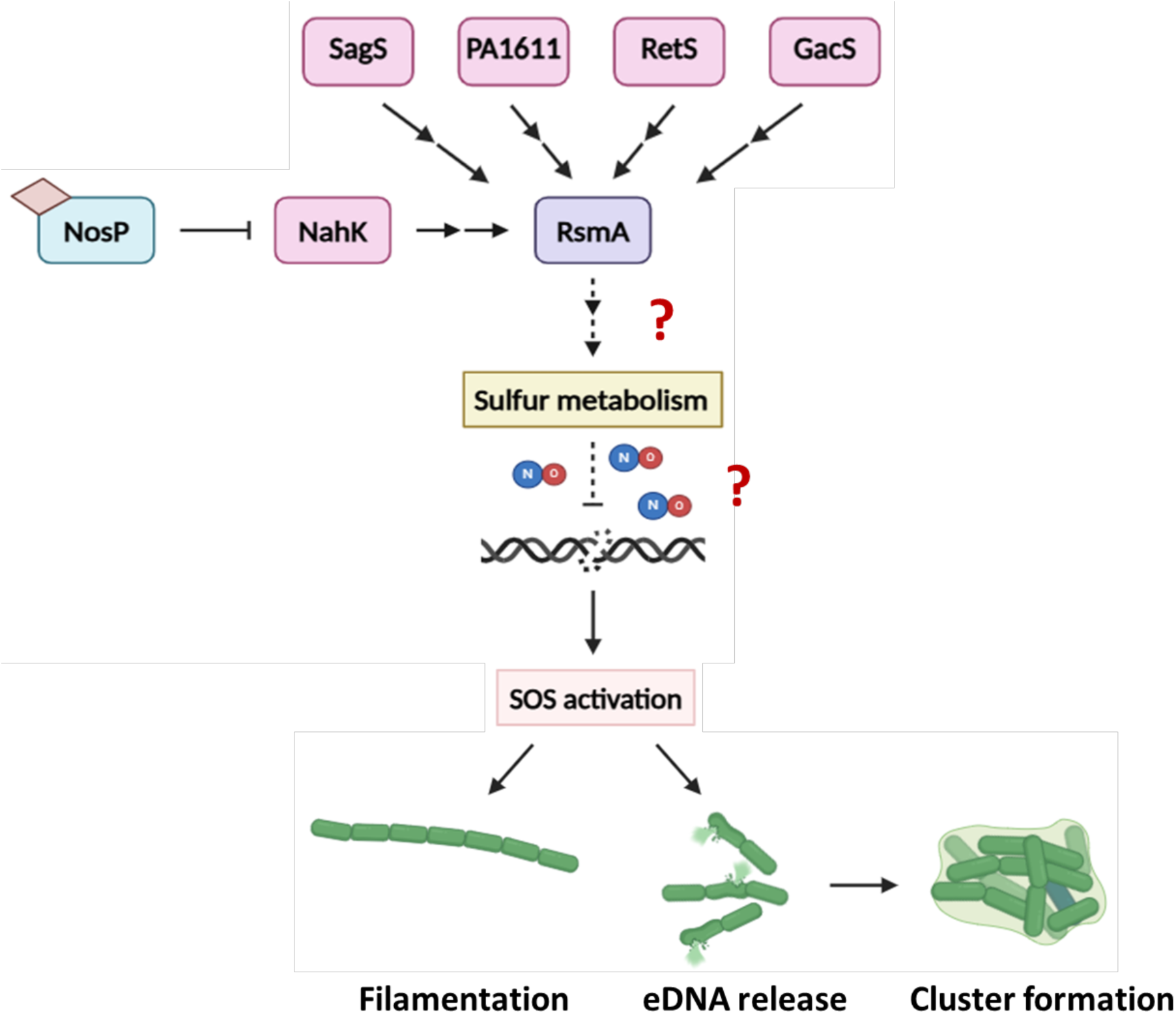
NahK prevents NO-mediated stress in *P. aeruginosa*. In *P. aeruginosa* PA14, NahK functions as one of five kinases that modulate the activity of the global post-transcriptional regulator RsmA. Active NahK ensures proper functioning of RsmA, which positively regulates sulfur metabolism genes through a yet unknown pathway. When NahK is inhibited, RsmA function is compromised, which decreases the transcription of sulfur metabolism genes. In minimal media lacking sufficient cellular sulfate pools, the presence of NO damages DNA and activates the SOS stress response, which ultimately leads to phenotypes such as cell filamentation and cell cluster formation due to eDNA released through cell lysis.

This work also aims to understand how the response of NahK to NO may be modulated by the growth conditions of the strain, especially in terms of the media used. *P. aeruginosa* is highly adaptable to various environments owing to its versatile metabolism (22–24). A previous study exploring the role of NahK in denitrification clearly shows the difference of PA14 WT strain to NO in aerobic and anaerobic conditions, with cell filamentation observed during anaerobic growth but not during aerobic growth (19). One of the key phenotypes of the Δ*nahK* strain is the production of the exotoxin pyocyanin due to mis-regulated quorum sensing and PQS accumulation, through inactivation of RsmA. Notably, pyocyanin production by *P. aeruginosa* PA14 has been shown to cause enhanced auto-poisoning and cell lysis of the producer cells in nutrient starved minimal media when compared to rich media (30). Similarly, biofilm formation in *P. aeruginosa* is more favored in media with high iron sources compared to media with depleted iron content (28, 29). Moreover, the role of the PAS domain in NahK is still unexplored and may serve a function based on the media components. Hence, in this work, the role of NahK in responding to NO is investigated in a minimal medium starved of nutrients such as iron and amino acids, to elucidate how these nutrients may affect the functioning of NahK specifically in its response to NO.

An NO-mediated SOS response has been observed in *P. aeruginosa* previously, albeit under different circumstances. PAO1 WT grown anaerobically not only caused cell filamentation due to NO accumulated during denitrification but also formed highly cohesive cell clumps. Affected cell division was found to be the cause of cell filamentation. Formation of cell clumps was linked to changes in membrane properties, not EPS components such as pel or psl, due to lack of upregulation of their regulatory genes (21). In light of our observations that cell filamentation due to NO-stress is accompanied by lysis-mediated eDNA release (**Fig. 5**), it is possible that the cell clumps observed previously were also formed due to eDNA.

Similarly, it has been reported that anaerobically grown PAO1 subjected to NO exposure through denitrification resulted in RecA mediated SOS response due to hindered DNA synthesis. This not only resulted in pyocin release but also in the formation of membrane vesicles (MVs), which are vesicles derived from the outer membranes of bacterial cells, and harbor proteins, DNA, and other cargo that enable microbial interaction and biofilm formation (65–68). MV formation was also observed during CPX triggered SOS response in PAO1, presumably due to membrane blebbing caused by a delay in cell division (69). Another method of MV formation is through explosive cell lysis, as observed in interstitial biofilms formed by *P. aeruginosa* PAK strain. Explosive cell lysis was observed in stress inducing conditions, due to activation of pyocin production and the concomitant lysis mediated by the endolysin Lys encoded in the pyocin gene cluster. Vesicularization of the cellular fragments generated through explosive cell lysis resulted in the formation of MVs containing cytosolic DNA and proteins (56). Our data depict significant upregulation of *lys*, pyocin genes, as well as *alpBCDE* genes involved in programmed cell death (**Fig. 6**). While it is not known whether cell lysis observed in our experiments is due to Lys-mediated explosive cell lysis or programmed cell death, or both pathways, there is enough evidence to suggest that MVs may be generated by NO treated Δ*nahK* strain as a result of SOS response. MVs not only enhance biofilm formation through interaction with eDNA but can also be cytotoxic to macrophages (67–69). It would be interesting to examine whether the Δ*nahK* strain exposed to macrophage-generated NO in a cell culture model also displays increased cytotoxicity due to stress mediated MV production.

Our research has shown that increased susceptibility of the Δ*nahK* strain to NO is due to decreased RsmA function in this strain (**Fig. 8**). RsmA, or its homolog CsrA, is a post-transcriptional regulatory protein well conserved across several bacterial species, controlling gene expression through repression of translation. This allows RsmA/CsrA to influence essential cellular processes in bacteria such as motility, virulence, biofilm formation, stress response and metabolism (13, 70–73). In *Escherichia coli*, nutrient limitation during growth in minimal media or during stationary phase of growth relieved CsrA repression of target mRNAs through the elevation of *csrB*/*C* transcription. Supplementation of amino acids in the form of tryptone or casamino acids significantly inhibited the transcription of *csrB*/*C*, thereby linking CsrA and nutrient starvation (74). Likewise, in *Vibrio cholerae*, the transcription of CsrA’s antagonistic srRNAs *csrB*/*C*/*D*, was higher in minimal media lacking essential amino acids such asparagine, arginine, glutamic acid and serine (75). In *Vibrio alginolyticus*, CsrA also significantly affected the genes involved in the metabolism of amino acids such as serine, alanine, proline, threonine and glutamine (76). More relevant to our studies, CsrA in *Erwinia amylovora* negatively regulated the transcription of sulfur metabolism genes (72). Although contrary to our results indicating positive regulation of sulfur metabolism genes by RsmA, this nevertheless supports the hypothesis that there may be a link between the two.

In *Legionella pneumophila*, *nahK* is encoded in the same operon as *nosP* and *narR*, which is a NahK associated response regulator that regulates biofilm formation through modulation of intracellular cyclic-di-GMP levels (77, 78). When *L. pneumophila* grows in a nutrient starved environment, the bacterium undergoes a lifestyle change from a replicative or exponential phase to a post exponential transmissive phase where the microbe may transmit to a new host. Interestingly, the *nosP*-*nahK*-*narR* operon was induced during the transition from exponential to post-exponential phase and during growth in minimal media, indicating that this operon was necessary for survival in low nutrient conditions. Notably, the shift from exponential to post-exponential phase is also facilitated in part by the alleviation of repression of CsrA targets, suggesting a link between CsrA, NahK, and nutrient starvation (78). While the molecular mechanism underlying NahK-mediated survival under nutrient starvation in *L. pneumophila* is currently unknown, it would be reasonable to postulate that such a regulatory pathway may also exist in a more metabolically evolved organism such as *P. aeruginosa*.

Our results suggest that NahK in *P. aeruginosa* may be involved in sulfur assimilation during conditions of sulfate starvation (SS), such as during growth in minimal media (**Fig. 10**). SS induced response in *P. aeruginosa* ensures the uptake and metabolism of alternate sulfur sources such as sulfate esters, glutathione and alkane sulfonates when preferred sulfur sources such as inorganic sulfate, thiocyanate or cysteine are depleted (60–62). This response is well studied in *E. coli*, where CysB, a primary regulator of cysteine biosynthesis, and Cbl, a CysB-like protein, regulates the expression of these genes. Depletion of cellular sulfate pools leads to the accumulation of cysteine precursor N-acetylserine, which acts as a co-inducer to activate CysB, leading to the upregulation of SS induced genes. Besides genes involved in uptake and metabolism of varied sulfate sources, SS was also linked to the biosynthesis and regulation of Cys and Met, as sulfur is an essential precursor for their biosynthesis. SS could also hinder the biosynthesis of Fe-S clusters as the *iscRSUA* operon, whose protein products ensure the incorporation of Fe-S clusters into proteins, was also upregulated. In *E. coli*, SS was also implicated in oxidative stress as it increased the transcription of *ahpCF* operon, which is normally activated in response to H_2_O_2_ stress by OxyR, an oxidative stress response regulator. Hence, SS may cause H_2_O_2_ accumulation, perhaps due to the depletion of cellular antioxidants such as cysteine and glutathione or because of impaired formation of Fe-S cluster or SH-containing enzymes involved in maintaining cellular redox balance (62). Similarly, in *P. fluorescens*, lack of sulfate in the growth medium triggered increased activity of antioxidant enzymes such as superoxide dismutase and catalase (79). In *P. aeruginosa*, unlike in *E. coli*, Cys and Met biosynthesis genes were not as significantly upregulated along with SS induced genes. However, there may still be a link between SS and oxidative stress, as *ahpC*, *katB*, *ohr* and *lsfA*, whose protein products are involved in combating oxidative stress, were all upregulated (80). According to previously published transcriptomic studies, *lsfA* is 5-fold downregulated in Δ*nahK* and Δ*rsmA* strains compared to WT, and we have shown that *lsfA* is positively regulated by RsmA in the qPCR studies reported here (15). These results suggest an additional reason why Δ*nahK* and Δ*rsmA* strains may be more susceptible to nitrosative stress than the WT strain.

It is not yet known why a significant number of sulfur metabolism genes are downregulated in the Δ*nahK* and Δ*rsmA* strains, compared to WT, when cultured in minimal media (15). Recent studies pertaining to CysB in *P. aeruginosa* show that it can regulate virulence through positive regulation of RetS, which is one of the five histidine kinases in the Gac multikinase network that regulate RsmA (81). RetS inhibits GacS, which leads to decreased transcription of *rsmY* and *rsmZ*, ultimately resulting in turning ON RsmA (14, 25, 82). A *cysB* deletion strain displayed increased expression of *rsmY*/*Z* and *gacA*, decreased expression of type III secretion system, reduced swarming motility and formed tower-like biofilms. These phenotypes are consistent with the loss of RetS mediated GacS repression, leading to RsmA being in its OFF state. However, CysB mediated regulation of RetS was not dependent on the availability of sulfur sources in the culturing media (81). Yet, we cannot rule out the possibility that in Δ*nahK,* when RsmA is OFF, there may be an autoregulatory feedback loop misregulating CysB and inadvertently interfering with sulfur metabolism. Consistent with this hypothesis, *cysB*, whose transcription is regulated by its own protein product, was significantly downregulated in Δ*nahK* and Δ*rsmA* strains compared to WT, albeit with a log_2_ fold change of less than −1 (15). This may also explain the misregulated denitrification and decreased biofilm formation of Δ*nahK* under anaerobic conditions compared to WT as sulfur metabolism is essential for anaerobic biofilm formation (19, 83). Transcriptomic studies of anaerobically grown PA14 biofilms revealed that 40 sulfur metabolism genes were significantly upregulated, presumably due to their role in the synthesis of Fe-S cofactors for denitrification enzymes such as Anr and Dnr. This was attributed to continued exposure to anaerobic environments rather than due to sulfate starvation (83). Hence, although the Δ*nahK* strain was grown anaerobically in rich LB media, optimum sulfur metabolism may have been necessary for sustainable function of denitrification reductases, the dearth of which would have resulted in decreased biofilm formation (19). However, down regulation of *cysB* in Δ*nahK* strain did not seem to affect *retS* transcription, as RetS still exerted its inhibitory effect on GacS in the Δ*nahK* strain (15). This further highlights the complex nature of the various interconnecting regulatory networks in *P. aeruginosa* and warrants additional studies to deconvolute a possible connection between NahK, RsmA, and CysB.

Taken together, our data contributes to expanding our knowledge of the different roles of NahK in biofilm regulation. Previous studies on NahK in *P. aeruginosa* revealed that Δ*nahK* strain is a pro-biofilm mutant; it forms robust three-dimensional biofilms, has decreased swarming motility and overproduces pyocyanin. Pyocyanin is not only a virulence factor but also facilitates respiration in micro-aerobic conditions and aids in the maintenance of biofilm architecture through interaction with eDNA (15, 16, 47). These phenotypes suggest that NahK may be inhibited within the deeper layers of a biofilm, presumably due to interaction with heme bound NosP (17). When NO is generated through denitrification, within the oxygen depleted deeper layers of a biofilm, NosP-NahK interaction may be altered due to NO ligation by NosP, leading to dispersal of a subpopulation of cells to find new surfaces for nourishment. Alternatively, or in addition, in a different sub-population of NahK-inhibited cells that are experiencing a nutrient starvation, exposure to NO may initiate the SOS response pathway to ensure survival. The resulting cell filamentation and lysis-mediated eDNA release of this subpopulation may be altruistic and benefit the surrounding cells at large by providing cytosolic nutrients. Interestingly, endogenous stress in biofilm microcolonies due to ROS, RNS, or nutrient starvation have been shown to cause cell lysis and dispersal, resulting in voids within the centers of microcolonies (32, 84, 85). The SOS response may also facilitate the production of MVs for QS and horizontal gene transfer, as well as induce antibiotic resistance, as the DNA damage repair occurring during SOS response is error prone and can induce favorable mutations. Similarly, RecA activation has been shown to increase genetic variants in a biofilm, presumably due to recombinatorial repair of DNA double strand breaks (86, 87). Further studies are necessary to investigate and elucidate a potential role for NahK in influencing cell death, antibiotic resistance and perhaps even persistence within *P. aeruginosa* biofilms, so that suitable targets in these pathways could be identified for developing new therapeutic strategies to treat *P. aeruginosa* biofilm associated infections.

## Materials and Methods

### Strains and growth conditions

All bacterial strains and plasmids used in this study are described in **Tables S1 and S2**. All strains were revived from glycerol stocks through overnight growth in LB media at 37℃ and 225 rpm. Overnight cultures were diluted 25-fold into 5 mL of fresh M9 minimal media (9 mM NaCl, 22 mM KH_2_PO_4_, 48 mM Na_2_HPO_4_, 19 mM NH_4_Cl, 1 mM MgSO_4_, 0.1 mM CaCl_2_) supplemented with 0.4% (w/v) glucose as the carbon source. The cultures were grown to an OD of 0.45, by shaking at 37℃ and 225 rpm, and then incubated with 500 μM of the iron chelator 2, 2’-bipyridyl. For experiments that required nitric oxide (NO) scavenging, 2 mM 2-(4-Carboxyphenyl)-4,4,5,5-tetramethylimidazoline-1-oxyl-3-oxide potassium salt, 2-(4-Carboxyphenyl)-4,5-dihydro-4,4,5,5-tetramethyl-1H-imidazol-1-yloxy-3-oxide potassium salt (PTIO) was added to the media at this stage. To further exhaust all the nutrients in the media, all the cultures were re-diluted 50-fold in fresh M9 minimal media and then grown overnight in the presence or absence of DETA NONOate. During this second overnight growth, wherever required, cultures were supplemented with 5 μM ferric chloride (FeCl_3_), 10 μM hemin, 0.5% (w/v) casamino acids (CAA), 0.025 μg/mL ciprofloxacin (CPX), 1 mM cysteine (Cys), 1 mM methionine (Met), 50 μM N-ethylmaleimide (NEM), 100 μM sodium nitrate (NaNO_3_), 1 μM M64, 4 mM Tiron or 50 μM serine hydroxamate (SHX). The amount of DETA NONOate added was determined by the volume of these second-generation minimal media cultures. 50 mL cultures were grown in 250 mL Erlenmeyer flasks in the presence or absence of 100 μM DETA NONOate whereas 5 mL cultures were grown in standard 5 mL capacity culture tubes, in the presence or absence of 50 μM DETA NONOate.

### Growth curves

Overnight cultures of PA14 strains were diluted in M9 minimal media and grown to an OD_600_ of 0.45, followed by 3 hours of iron chelation and re-inoculation into fresh M9 minimal media. To test the effect of NO on the growth of WT and Δ*nahK* strains, 5 mL cultures were set up in the presence or absence of 100 μM DETA NONOate, which releases approximately 100 nM of NO gas. This concentration of DETA NONOate was used in order to mimic the experimental conditions previously employed to evaluate the response of WT and Δ*nahK* strains to NO under aerobic growth in rich media and to determine if this response was altered in a nutrient starved media (19). However, hardly any growth was observed in the Δ*nahK* strain and it was hypothesized that the smaller headspace in the culture tubes coupled with increased susceptibility to NO in minimal media may have resulted in this lack of growth. Hence, we repeated the experiment in 50 mL minimal media in the presence and absence of 100 μM DETA NONOate. To test the effect of iron and amino acids on the growth of NO-treated Δ*nahK* strain, 5 mL cultures were set up in the presence or absence of 50 μM DETA NONOate in minimal media supplemented with or without 5 μM FeCl_3_, 10 μM hemin or 0.5% (w/v) CAA. Cultures were then grown overnight for 24 hours at 37℃ and with shaking at 225 rpm. The OD_600_ of each culture was measured using an optical density meter every hour for the first 16 hours, then again at t=20 hours and t=24 hours.

### Construction of deletion and reporter strains

All deletion strains were constructed using previously described methods and all primers used are listed in **Table S3** (88). Briefly, the upstream and downstream flanking regions (approximately 900 bp) of the gene of interest were amplified through PCR using PA14 genomic DNA as the template. Here, the genes of interest were PA14_54430 (*algU* locus tag in PA14), PA14_29640 (*fhp* locus tag in PA14) and the first 1377 bp ( or 459 amino acids) of PA14_38970 (*nahK* locus tag in PA14), which encodes for the PAS domain of NahK. The PCR amplified flanking regions were joined together by overlap extension PCR and the resulting product was cloned into the allelic exchange vector pEX18(Gm). This generated 3 plasmids; pEX18(Gm)_Δ*algU*, pEX18(Gm)_Δ*fhp* and pEX18(Gm)_ΔPAS *nahK*. These plasmids were then electroporated into the *E. coli* mating strain SM10 and introduced into PA14 strains through biparental mating. pEX18(Gm)_Δ*fhp* and pEX18(Gm)_ΔPAS *nahK* were introduced into PA14 WT strain while pEX18(Gm)_Δ*algU* was introduced into PA14 Δ*nahK* strain to generate a double deletion mutant strain. The colonies that took in the plasmid were selected on *Pseudomonas* isolation agar (PIA) plates containing 60-100 μg/mL gentamicin and confirmed by colony PCR. For making markerless deletions, the confirmed colonies were streaked on LB agar plates containing 10-15% sucrose and lacking NaCl, for counterselection, and the resulting clones were confirmed by colony PCR. To construct the P*fhp*-*lacZ* reporter, the promoter region of *fhp* (approximately 700 bp) was PCR amplified from PA14 genomic DNA and cloned into a mini-CTX-*lacZ*(Tet) vector. The resulting plasmid was electroporated into *E. coli* SM10 cells and introduced into PA14 WT and Δ*nosP*Δ*nahK* stains by the biparental mating method described above. Colonies that took in the plasmid were selected on PIA plates containing 200 μg/mL tetracycline and confirmed by colony PCR.

### Cell morphology imaging and quantification

Day cultures of PA14 strains in M9 minimal media were subjected to iron chelation and inoculated into 5 mL of fresh M9 minimal media. The cultures were then grown overnight for 16 hours at 37℃ with shaking at 225 rpm, in the presence or absence of 50 μM DETA NONOate, with or without other additives in the media, as required for the experiment. 500 to 1000 μL of the stationary phase cultures were centrifuged for 1 minute at 14,000 rpm and cell pellets were collected after discarding the supernatant. The fixing solution 4% paraformaldehyde was used to redissolve the cells pellets and 5 μL of the resulting solution was pipetted onto a glass slide, heat fixed for 10 seconds and then covered by a glass coverslip. Microscopy images were recorded using a Zeiss Axio Vert.A1 inverted transmitted light microscope equipped with a Lumencor Sola Light Engine light source and a Zeiss A-Plan 40x N.A. 0.55 objective. The images were scaled using ImageJ software and the cell lengths of randomly selected cells were determined using the select line tool on the software. The experiment was carried out in both small scale 5 mL cultures and large scale 50 mL cultures, with 50 μM and 100 μM DETA NONOate respectively, to evaluate whether the same extent of stress was induced in both cultures. Both small scale (50 μM DETA NONOate in 5 mL culture) and large scale (100 μM DETA NONOate in 50 mL culture) NO-treated cultures displayed similar cell filamentation phenotypes, suggesting that the higher concentration of NO in the large-scale cultures may be compensated for by the increased headspace in these cultures (**Fig. S12**).

### RNA extraction and quantitative reverse transcription PCR (qPCR)

Day cultures of PA14 strains in M9 minimal media were subjected to iron chelation and inoculated into 50 mL of fresh M9 minimal media. The cultures were then grown overnight for 16 hours at 37℃ with shaking at 225 rpm, in the presence or absence of 100 μM DETA NONOate, with or without other additives such as CAA in the media, as required for the experiment. 2 mL of the stationary phase cultures were treated with 4 mL of the RNAprotect Bacteria Reagent (Qiagen) and were either frozen at −80℃ or used directly. The total cellular RNA was isolated from the cell pellet using the RNeasy Mini Kit (Qiagen) and its concentration and purity were determined using Nanodrop and 1% agarose gel. 1 μg of purified RNA from each sample was used to make complementary DNA (cDNA) using the Maxima First Strand cDNA Synthesis Kit (Thermo Scientific). Each cDNA stock was diluted 3-fold for the final qPCR reaction, which also comprised 0.3 μM forward and reverse primers (**Table S3**) and 1x Maxima SYBR green qPCR master mix (Thermo Scientific). PCR was carried out on the resulting 10 μL reaction mixture, on a LightCycler 480 thermocycler, with the cycle parameters set to 10 min initial denaturation at 95℃, followed by 40 cycles of 15 s denaturation at 95℃, 30 s annealing at 60℃ and 30 s extension at 72℃. The Ct values of the target genes and the housekeeping gene *proC* were obtained for all the cultures tested and the fold change of each gene in all the tested conditions was calculated relative to the untreated WT culture using the 2^(-ΔΔCt) method.

### Quantification of extracellular DNA

Day cultures of PA14 strains in M9 minimal media were subjected to iron chelation prior to inoculation into 5 mL of fresh M9 minimal media. These cultures were grown overnight for 16 hours at 37℃ with shaking at 225 rpm, in the presence or absence of 50 μM DETA NONOate, with or without other supplements in the media, as required for the experiment. Stationary phase cultures were diluted in M9 minimal media to an OD_600_ of around 0.1 and 500 μL of each culture was transferred to one well of a 4-well Nunc Lab-Tek II Chamber Slide (Thermo Scientific). A rapid attachment assay was carried out wherein each culture was incubated in the chamber slide for 15 minutes at 37℃ to allow for all the live cells to attach using their active pili machinery. After the incubation period, the cultures were discarded from the wells and each well was rinsed with 500 μL of fresh M9 minimal media twice. This was done to remove dead cells that would have failed to attach to the wells in such a short period of time due to lack of an active pili machinery. For visualization of extracellular DNA (eDNA), the red-fluorescent membrane impermeable nucleic acid stain propidium iodide (PI) was used. PI stain was diluted in M9 minimal media to a final concentration of 10 μM and incubated with the samples in each well for 5 minutes at room temperature, after which the PI solution was discarded and replaced by fresh M9 minimal media. This was followed by imaging using the Zeiss Axio Vert.A1inverted transmitted light microscope equipped with a Lumencor Sola Light Engine light source, a Zeiss A-Plan 40x N.A. 0.55 objective and a Texas Red fluorescence filter (excitation wavelength - 560 nm, emission wavelength - 630 nm). Multiple brightfield and red fluorescence microscopy images were recorded for each sample, by scanning through each well. Samples were considered to have accumulated eDNA sites if red fluorescence was observed in the fluorescence microscopy images and the corresponding brightfield microscopy images displayed cells aggregated together due to the sticky nature of eDNA. The extent of eDNA accumulation in each strain was then quantified by manually counting the number of eDNA sites in 30 random fields of view (FoV) across 3 biological replicates.

### RNA sequencing

Overnight cultures of PA14 strains were diluted into 5 mL of M9 minimal media and grown to an OD_600_ of 0.45 followed by 3 hours of iron chelation and re-inoculation into 50 mL of fresh M9 minimal media. The 50 mL cultures were either supplemented with or without 100 μM DETA NONOate and grown overnight for 16 hours at 37℃ with shaking at 225 rpm. All the conditions included 2 biological replicates and the total cellular RNA from all the cultures were isolated as described previously for qPCR analysis. The quality and concentration of the extracted RNA were then analyzed using a NanoDrop and a Bioanalyzer 2400. RNA sequencing was performed using an Illumina NextSeq 550, on a mid output 150 cycle flow cell kit assembly, at the Stony Brook University Genomics and Bioinformatics Core facility. Illumina generated sequencing reads were processed and mapped to the *Pseudomonas aeruginosa* UCBPP-PA14 genome sequence from the *P. aeruginosa* genome database (https://pseudomonas.com/) manually. The relative expression levels of all genes were reported as log_2_ fold change and all genes reported with *p-*value > 0.05 were disregarded from further analysis.

### β-galactosidase activity assay

Overnight cultures of PA14 strains, electroporated with P*fhp*-lacZ reporter plasmid, were diluted into 5 mL of M9 minimal media and grown to an OD_600_ of 0.45 followed by 3 hours of iron chelation and re-inoculation into 50 mL of fresh M9 minimal media. After overnight growth in media supplemented with or without 100 μM DETA NONOate, 20 μL of each culture was mixed with 80 μL of permeabilization solution (100 mM Na_2_HPO_4_, 20 mM KCl, 2 mM MgSO_4_, 0.8 mg/mL hexadecyltrimethylammonium bromide, 0.4 mg/mL sodium deoxycholate, 5.4 μL/mL β-mercaptoethanol) followed by the addition of 600 μL of substrate solution (60 mM Na_2_HPO_4_, 40 mM NaH_2_PO_4_, 1 mg/mL o-nitrophenyl-β-D-galactoside, 2.7 μL/mL β-mercaptoethanol). The reaction was incubated at 37℃ for 47 min and then quenched using 700 μL of stop solution (1 M Na_2_CO_3_). 100 μL of the quenched reactions were then transferred into a 96-well plate and the absorbance at 420 nm was recorded using an Agilent BioTek Synergy H1 plate reader. The Miller Units were calculated using the equation (1000 x Abs_420_)/[Abs_600_ x (vol. 0.01 mL) x (reaction time in min)].

## Data Availability

All data supporting the findings of this study are available from the corresponding author Elizabeth Boon upon request. All constructs and cell lines associated with this study are freely available upon request.

## Acknowledgements

We thank Danielle Guercio (Department of Molecular and Cellular Biology, Stony Brook University) for helpful discussion of the manuscript. We thank Petr Dao (Department of Chemistry, Stony Brook University) for assisting with the generation of *algU* deletion strains. We thank Scott Laughlin (Department of Chemistry, Stony Brook University) and members of the Laughlin lab for providing training and access to the inverted transmitted light microscope utilized for imaging bacteria. We thank Jessica Seeliger (Department of Pharmacological Sciences, Stony Brook University) and members of the Seeliger lab for providing training and access to the thermocycler utilized for qPCR analysis. We thank the Renaissance School of Medicine Genomics Core Facility, Stony Brook University, for supporting the RNA sequencing studies. This study was supported by the National Institutes of Health (NIH) grant R01GM118894 provided to E.M.B.

## Author Contributions

S.A. - conceptualization, data curation, formal analysis, investigation, methodology, validation, visualization, writing – original draft, writing – review and editing; J.F. - conceptualization, data curation, formal analysis, investigation, methodology, validation, visualization, writing – review and editing; E.M.B - conceptualization, formal analysis, funding acquisition, project administration, resources, supervision, visualization, writing – review and editing.

## Supplemental Material

Supplemental tables and figures. Tables S1 to S9; Figures S1 to S12

